# Functional screen of Inflammatory bowel disease genes reveals key epithelial functions

**DOI:** 10.1101/2021.10.15.464566

**Authors:** Jessy Carol Ntunzwenimana, Gabrielle Boucher, Jean Paquette, Hugues Gosselin, Azadeh Alikashani, Nicolas Morin, Claudine Beauchamp, Louise Thauvette, Marie-Ève Rivard, Frédérique Dupuis, Sonia Deschenes, Sylvain Foisy, Frédéric Latour, Geneviève Lavallée, Mark J. Daly, Ramnik J. Xavier, the iGenoMed Consortium, Guy Charron, Philippe Goyette, John D. Rioux

## Abstract

**Background:** Genetic studies have been tremendously successful in identifying genomic regions associated with a wide variety of phenotypes, although the success of these studies in identifying causal genes, their variants, and their functional impacts have been more limited.

**Methods:** We identified 145 genes from IBD-associated genomic loci having endogenous expression within the intestinal epithelial cell compartment. We evaluated the impact of lentiviral transfer of the open reading frame (ORF) of these IBD genes into the HT-29 intestinal epithelial cell line via transcriptomic analyses. Comparing the genes whose expression was modulated by each ORF, as well as the functions enriched within these gene lists, identified ORFs with shared impacts and their putative disease-relevant biological functions.

**Results:** Analysis of the transcriptomic data for cell lines expressing the ORFs for known causal genes such as HNF4a, IFIH1 and SMAD3 identified functions consistent for what is known for these genes. These analyses also identified two major cluster of genes: Cluster 1 contained the known IBD causal genes IFIH1, SBNO2, NFKB1 and NOD2, as well as genes from other IBD loci (ZFP36L1, IRF1, GIGYF1, OTUD3, AIRE and PITX1), whereas Cluster 2 contained the known causal gene KSR1 and implicated DUSP16 from another IBD locus. Our analyses highlight how multiple IBD gene candidates impact on epithelial structure and function, including the protection of the mucosa from intestinal microbiota, and demonstrate that DUSP16, acts a regulator of MAPK activity and contributes to mucosal defense, in part via its regulation of the polymeric immunoglobulin receptor, involved in the protection of the intestinal mucosa from enteric microbiota.

**Conclusions:** This functional screen, based on expressing IBD genes within an appropriate cellular context, in this instance intestinal epithelial cells, resulted in changes to the cell’s transcriptome that are relevant to their endogenous biological function(s). This not only helped in identifying likely causal genes within genetic loci but also provided insight into their biological functions. Furthermore, this work has highlighted the central role of intestinal epithelial cells in IBD pathophysiology, providing a scientific rationale for a drug development strategy that targets epithelial functions in addition to the current therapies targeting immune functions.

## BACKGROUND

The inflammatory bowel diseases (IBD) are characterized by chronic relapsing inflammation of the gastrointestinal tract. Crohn’s disease (CD) (MIM 266600) and Ulcerative colitis (UC) (MIM 191390) are the two main subtypes of IBD. IBD are complex diseases, and as such are influenced by multiple contributing genetic and non-genetic risk factors. Genome-wide association studies (GWAS) have identified and validated over 200 genomic regions associated with IBD, with the majority being shared by CD and UC (1, 2). However, GWAS alone have only infrequently directly identified the actual causal gene or variant(s) in any given locus (3, 4). In order to address this challenge, high density genotyping and targeted sequencing studies of a subset of these IBD loci have been performed, successfully identifying candidate causal variants that implicate specific genes within some loci as being causal (5–8). Functional studies have validated many of these (6, 9–12). More recently, whole exome sequencing (WES) and whole genome sequencing (WGS) studies have identified additional loci as well as likely causal genes within these loci and previously known loci (13–15). Taken together, these studies have identified the causal gene, with one or more causal variants per gene, for two dozen of the validated IBD genetic loci. Importantly, these studies have enabled the identification of biological functions that are key to the development of IBD, such as microbial recognition, autophagy, cytokine signaling, and intestinal epithelial barrier function (3, 4, 12, 16–18).

Given that the causal gene and their variants remain to be identified for most known loci, we sought to develop a complementary approach for analysing known GWAS loci. This approach is based on the observations that (1) most causal variants in IBD have an impact on gene expression as opposed to gene functions (19–21) , (2) a gene’s function is believed to be influenced by its cellular context, and (3) the intestinal epithelium plays a key role in the development of IBD. Indeed, we set out to expand our understanding of biological functions relevant to IBD by performing an expression-based functional screen of genes from IBD loci in a human cell line commonly used as a model of intestinal epithelium.

Specifically, we created cell lines via lentiviral transfer of open reading frame (ORF) of 145 genes from validated IBD loci that had significant endogenous expression within human intestinal tissues and/or cell lines into the human colon adenocarcinoma HT-29 cell line. The biological impact of the expressing these different ORFs was assessed using transcriptomics. Initial analyses of this data demonstrated that the observed effects on the transcriptome were robust, reproducible, and were biologically relevant. This transcriptomic data was then analyzed to identify shared or opposite effects between IBD genes, including genes known to be causal. Overall, this approach led to the identification of clusters of genes with shared effects on the epithelial cell’s transcriptome. These results suggest that IBD genes impact a broad set of functions important to the intestinal epithelium barrier function and its contribution to host anti-microbial defense.

## METHODS

### Selection of candidate genes from IBD-associated regions

In order to identify the IBD gene candidates to test in our transcriptome-based screen for IBD gene functions, we focused on the 163 IBD-associated loci identified by the International IBD Genetics Consortium (1). First, we selected all genes found within a linkage disequilibrium (LD) region defined around each index single-nucleotide polymorphism (SNP; r_2_≥0.8 using the SNAP (22) online tool on the pilot 1K genome dataset (GRCh37/hg19)). Second, for regions where no genes were identified within the LD region, we selected the closest gene on either side of the index SNPs. For IBD-associated regions where the causal gene was already known (NOD2, IL23R, ATG16L1, IRGM, MST1, CARD9), only the causal gene was considered. From the combined list of genes identified in this manner, we removed the major histocompatibility complex (MHC) region and HOX gene clusters, as well as genes encoding known secreted proteins (cytokines, chemokines, etc) and receptors (identified through bioinformatic analysis), as these classes of proteins are unlikely to yield any information in an ORF-expression based screen without their required ligand or receptor. Finally, we removed from this list any gene that did not have detectable RNA expression in human intestinal tissues (small intestine, colon) or cell lines (HT-29, HCT-15, Caco-2) from our profiling experiment described below, although three known IBD causal genes were kept regardless of their lack in epithelial expression. In total, 169 genes from IBD-associated loci were selected for our expression-based screen in human colonic epithelial cells (**Additional file 1: Table S1, Additional file 2: Fig.S1**).

### Cloning of IBD Open Reading Frames (IBD-ORFs)

For the ***screening stage*** of this study, we set-out to clone the ORFs for all the IBD gene candidates selected above into our modified GATEWAY® compatible polycistronic lentiviral expression vector, pLVX-EF1a-IRES-PURO/eGFP expressing an enhanced green fluorescent protein (eGFP) and puromycin N-acetyl-transferase fusion protein, to use for expression in the HT-29 colonic epithelial cell line model. First, we queried the Ultimate_TM_ ORF Lite Clone Collection library (Invitrogen,Canada) for the presence of pEntry plasmid clones containing the longest validated isoform of each ORF, as defined by the NCBI’s Consensus CDS (CCDS) project. The cloned ORF sequences (supplied with the library) were then screened for point mutations affecting the primary amino acid sequence of the protein. In the cases where a SNP was found affecting the amino acid sequence, we only retained the cloned ORF if it contained the common allele. For any gene candidate either absent from the library, showing only smaller ORF isoforms, or carrying changes to the accepted primary sequence of the protein, a full-length codon-optimized attB-flanked sequence was synthesized using the Invitrogen GeneArt gene synthesis service (Invitrogen, Canada) and inserted into pDONR vector using BP reaction. All ORFs were then cloned into the pLVX-EF1a-IRES-PURO/eGFP vector using LR reaction, downstream of the constitutive promoter and upstream of an IRES-controlled Puromycin-eGFP hybrid reporter gene, so that transfection, transduction and expression could be monitored by fluorescence. Cloning was performed using the GATEWAY® cloning system (ThermoFisher Scientific), and cloned ORFs were fully sequenced to rule out any point mutations or cDNA rearrangements impacting on the primary protein sequence. A total of 10 failed to clone in our expression vector and therefore were dropped from the analysis.

In addition to the ORFs for the IBD gene candidates identified within IBD-associated loci, we also cloned three IBD-associated non-synonymous coding variants of the IFIH1 gene (described in text) using the codon-optimized ORF sequence (GeneArt String Synthesis and GeneOptimizer technology, Invitrogen). Mutated IFIH1 ORF sequences were synthesized and cloned as described above and were treated moving forward as any of the other IBD gene candidate ORFs in this study.

### Cell lines and culture conditions

The HEK 293T/17 (ATCC CRL-11268) cell line used for lentivirus production was cultured in DMEM, 10% FBS and used at 50-80% confluency for transfection of lentiviral vector and packaging plasmids as described in Production of lentiviral stocks. The colorectal adenocarcinoma cell line HT-29 (ATCCHTB-38) was maintained at confluence between 20% and 80% at all times in McCoy’s 5A (Wisent, 317-010CL) 10% FBS. The HCT-15 (ATCC CCL-225) cell line was cultured in RPMI-1640 (Gibco, 61870-036), 10% FBS and also maintained sub-confluent. Typically, all cell lines were maintained at confluence between 20% and 80% at all time in a 5% CO_2_ humidified atmosphere at 37C. Cells were passaged with incubation in TRYPLE (ThermoFisher Scientific) at 37C for 9 min and then diluted in order get a confluency of 20%. All experiments described herein were performed in cell at passages below 25.

### Production of lentiviral stocks

Lentiviral particle stocks were produced for all cloned IBD gene candidate ORFs. Lentiviral expression vectors were purified with GenElute HP Plasmid Maxiprep Kit (Sigma, St-Louis, MO) and only plasmid preparations with absorbance (260/280) ratios 1.8 to 2.0 were used for transfection. Each ORF cloned into the pLVX-EF1a-IRES-PURO/eGFP lentiviral expression vector was added to lentiviral packaging and envelope vectors (Sigma-MISSION-gag-pol and the envelope Sigma-MISSION-VSV-G (Cat: CST-DNA Sigma, St-Louis, MO)) in a 2:2:1 ratio and co-transfected into HEK-293T packaging cells via the calcium phosphate precipitation method according to the Open-Biosystem protocol (as described in (23)). Following an incubation of 7-8 hours in DMEM supplemented with 10 % fetal bovine serum (FBS), 1% L-Glutamine 2 mM, 0.1% Pen-Strep at 37°C and 5% CO_2_ overnight at 37°C under 5% CO_2,_ the culture media was replaced with fresh culture media and the cells incubated at 37°C and 5% CO_2_ for an additional 40 hours. Lentivirus-containing medium was then harvested, cell debris pelleted, and viral particles were concentrated using the Lenti-X Concentrator (Takara Bio) and resuspended with McCoy’s 5A (Wisent) in 1/10 of the original volume (10X concentration). The lentiviral stock was divided in 100 ul single-use aliquots and stored at -80°C.

All pLVX-EF1a-IRES-PURO/eGFP lentiviral stocks containing the different IBD gene candidate ORFs were titrated on HT-29 cells (ATCC HTB-38). Proliferative HT-29 cells, grown in McCoy’s 5A (Wisent Cat: 317-100-CL) supplemented with 10 % FBS, 1% L-Glutamine 2 mM, 1% Penicillin-Streptomycin (Pen-Strep) at 37°C under 5% CO_2_, were transduced with 25ul of a 5-fold serial dilution (in McCoy’s 5A without serum) of each viral stock along with Polybrene (Sigma, Cat: H-9268) at a final concentration of 8ug/ml to improve transduction efficiency. Cells were then centrifuged at 1200 X g for 30 min at 37 °C to increase transduction efficiency and incubated for 4 hours before media was changed and cells cultured for an additional 48 hours. Independent eGFP-expressing cells/colonies were counted using an IN Cell 6000 confocal microscope (GE Healthcare, Marlborough, MA), average background fluorescent cell counts of non-infected cells subtracted and transduction values were converted in Transducing units per ml (TU/ml). All procedures were done in accordance with enhanced BSL-2 (biosecurity level-2) containment guidelines.

### Lentiviral transduction, tissue culture and antibiotic selection in HT-29 cells

Proliferative HT-29 cells at 50% confluency in 12 well plate (Corning) were transduced in triplicate with viral stock at a (multiplicity of infection) MOI of 40-100 with 8µg/ml of polybrene (Sigma, Cat: H-9268). Twenty-four hours post-transduction, media was changed and 3ug/ml puromycin (Millipore Sigma) was added. After 3 days of puromycin selection, the percentage of eGFP-expressing cells/colonies were counted using an IN Cell 6000 confocal microscope (GE Healthcare, Marlborough, MA) and any well presenting less than 5% green fluorescent protein (GFP) positive cells was eliminated. The appearance and confluence of the cultures were recorded daily, and cells were grown for an average of 8 days (with a range of 5-27 days) to select successfully transduced cells and reach confluence before RNA extraction. Transduction of the whole set of IBD gene candidate ORFs was performed in batches of about 15 ORFs, each done in triplicate, and including an empty vector control. Some ORFs were repeated between batches for quality control and validation purposes. A total of 12 ORFs failed to transduce into HT-29 cells and were dropped from the analysis.

### Transcriptomic analyses

RNA was extracted from transduced HT-29 cultures using the RNeasy Plus Mini kit (#74036, Qiagen Inc Canada) according to manufacturer’s protocol. The RNA samples were quantified, and quality controlled using an Agilent RNA 6000 Nano kit (Agilent, Cat No./ID 5067-1511) on 2100 Bioanalyzer system. The samples with RNA Integrity Number (RIN) below 8 were discarded; RIN values were routinely in the range of 9.2-10. Expression profiling of the RNA was performed using two expression arrays: a custom-built targeted Agilent array and an Illumina whole genome array. Specifically, the Agilent iGenex v.2 gene expression array (Agilent, Santa Clara, CA) is a custom-designed array with 31,344 probes targeting all exons of 2099 genes, including genes associated with immune-mediated and autoimmune diseases, markers of epithelial and immune differentiation and function, as well as housekeeping genes, whereas the Illumina Human HT-12 v4 beadchip (Illumina, San Diego, CA) is a genome-wide array targeting about 31,000 genes (with on average 2 probes per gene) with more than 47,000 probes. RNA sample labeling for the Agilent array was done with the Agilent One-Color Microarray-Based Exon Analysis Low Input Quick Amp WT Labeling kit and the arrays were processed on the Montreal heart institute (MHI) Integrative Biology Platform’s Agilent system (Agilent Microarray Hybridization oven and SureScan Microarray scanner) in batches of ∼15 ORFs in triplicate (∼45 samples), with triplicates randomized on different arrays. Some RNA samples were repeated between batches of arrays for quality control and evaluation of batch effect correction. Two of the three replicates for each ORF were randomly selected to be evaluated on the genome-wide Illumina array. Illumina arrays were processed at the Génome Quebec / McGill Innovation Centre in 5 batches, with replicates randomized between processing dates and arrays, with some RNA samples repeated. We have uploaded the raw expression data from the custom Agilent and the whole genome Illumina Human HT-12 v4 beadchip arrays onto GEO (accession numbers XXX (24) and YYY (25) respectively).

### Validation of IBD gene candidate ORF RNA expression

Expression of target ORFs was validated directly from the expression arrays transcriptomic data when possible. In order to validate the expression of ORFs that were not detectable on the expression arrays, due to the absence of functional probes within available exons, probes located only in untranslated region (UTR) sequences or codon-optimized ORF sequences, we developed an alternative end-point PCR approach. Briefly, a common forward primer located within the pLVX plasmid sequences 5’ to the ORF insertion site along with a reverse ORF-specific primer (designed using Primer 3Web) were used for end-point PCR **(Additional file 1: Table S2)**. Levels of eGFP RNA produced from the pLVX-EF1a-ORF-IRES-PURO/eGFP plasmid was also used to validate expression levels from the transduced plasmids; these were found to be within a similar expression range for all transduced ORFs (data not shown).

All ORF-specific PCR primer sequences used in this validation are shown in **Additional file 1: Table S3** and primers were obtained from Millipore Sigma unless mentioned otherwise from Integrated DNA Technologies. Total RNA preparations(1ug) from ORF transduced cell lines were reversed transcribed using the High Capacity cDNA RT kit (ThermoFisher Scientific) and end-point PCRs were performed following 34 cycles (94 C 30 sec, 58 C 30 sec, 72 C 1 min) using the HotStarTaq Plus DNA polymerase (Qiagen, Canada) according to manufacturer’s protocol.

### Description of samples

Overall, for the transcriptome-based screen (**Additional file 2: Figure S1**), we successfully cloned the ORFs, transduced in HT-29 in triplicate, generated stable cultures, and isolated RNA for 147 out of the 169 genes selected from IBD-associated loci for a total of 441 RNA samples (see **Additional file 1: Table S1** for details on the different ORFs transduced). All these samples were included in the targeted (Agilent) transcriptomic analysis and two of the three samples from each ORF were included in the genomewide (Illumina) transcriptomic analysis (see above). Two of the ORFs tested did not show over-expression in our cell model were removed from downstream analytical steps (see **Additional file 1: Table S4** for details on the expression levels of the different ORFs transduced). For replication purposes, additional transductions (n=3) were performed for 9 of these genes (27 RNA samples) (**Additional file 2: Figure S1**). For functional validation experiments, we transduced lentiviruses (n=6) containing the wild-type IFIH1 ORF, as ORFs carrying 3 IBD-associated variants (see results section for description), as well as additional transductions (n=3) for the DUSP16 and KSR1 ORFs (**Additional file 2: Figure S1**).

### Expression profiling of IBD gene candidates

Gene expression profiling of 297 genes from IBD-associated loci was performed across a panel of different RNAs from human tissues (n=1, purchased from Clontech Laboratories) and from different immortalized intestinal and immune cell lines (n=3, cell lines were obtained from ATCC and grown in-house, RNA was isolated using the RNeasy Plus Mini kit (#74036, Qiagen Inc Canada) according to manufacturer’s protocol) using our custom-made Agilent iGenex v.2 gene expression array (see above). An expression value was obtained for each gene in each replicate by calculating the geometric mean of all probes within the gene, followed by a median normalization across all genes on the array. A geometric mean was calculated from at least 3 independent measurements for each tissue. Given that only one sample was available for tissues, technical replicates were used.

### Analysis of Agilent chip data from HT-29 screen

We used Limma v3.26.9 and sva v3.18.0 packages (Bioconductor v3.1-3.2) (26, 27) to load the fluorescence intensity files and handle the data into R V3.2.0. Arrays failing Agilent quality control according to the Agilent Feature Extraction software were flagged. Manual quality control of each array was also performed. Arrays with high background signals, too low or saturated fluorescence intensities, poor recovery of spiked-in standards or non-uniform signal (as detected by Agilent quality metrics and/or visual inspection of the image) were removed. Each array was also aligned to the median of the whole batch and those showing bad alignment of the positive, negative or spiked in controls were removed. Each sample was then investigated to confirm the identity of the ORF that was expressed. Strong outliers not explained by biology were also removed. All removed samples were repeated in the next batch. Quality control was performed within each batch of arrays.

After stringent quality control, fluorescence signal was processed to remove local background (bakgroundCorrect, method “normexp”) and then normalized using cyclic loess algorithm. Batch effects were then corrected using ComBat (**28**) for known batches and sva for unknown systematic artefacts (both from sva library v3.18.0). Given the experiment was performed in three parts, the batches from each part were combined first and then combined together. To reduce impact of residual background, noise and batch effects on very low expression values, the fluorescence intensity values were truncated at 8, and translated down by the same amount. Duplicated probes were combined, and repeated samples removed. The truncating threshold was determined based on the distribution of negative control probes and detected vs non-detected signals.

### Analysis of Illumina chip data from HT-29 screen

Beadarray v2.20.1 and illuminaHuman4.db v1.24.0 packages (R Bioconductor v3.0) (29) were used to load and annotate (readBeadSummary and add FeatureData) the fluorescence intensity data into R-3.2.0. After normalization of the array data (normaliseIllumina, method=”neqc”), data was converted and handled using Limma package v3.26.9 (Bioconductor v3.0). Probes annotated as “Bad” or “No match” were removed. Manual quality control of each array was performed. Arrays with high background signals, too low, saturated or uneven fluorescence intensities were removed. Each array was also aligned to the median of the whole batch and those showing bad alignment of the positive, negative or housekeeping genes controls were removed. Strong outliers not explained by biology were also removed. Removed samples were repeated in the next batch. Quality control was performed within each batch of arrays.

Batch effects were then corrected using ComBat for known batches and sva v3.18.0 for unknown systematic artefacts. Given the chips were performed in five batches, each with at least two processing date, the dates from each batch were combined first before merging the batches. To reduce impact of residual noise or batch effects on very low expression values, the fluorescence intensity values were truncated at 2^3.5, and translated down by the same amount.

### HIT definition

Given that we expect expression of the different ORFs to have distinct effects on the expression pattern of different genes in the transcriptome, we leveraged the information from the complete dataset to generate the baseline distribution of expression for all genes expressed in the HT-29 cell line under the culture conditions used in this project. After log_2_ transformation of the expression data, the median expression measured by each probe over all samples was used as the baseline, while the median absolute deviation (MAD) was used to define expected range of variability. This information was summarized as a Z score computed as: 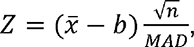 where *n* is the number of replicates (*n*=3), 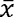 is the average from the replicates and 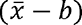 is the difference from the baseline. Probes with |Z| > 4 were considered outside expected range of variation. For each expressed ORF, a probe was defined as a HIT if the deviation 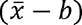 was larger than 1, which is equivalent to a fold effect greater than 2 on the original scale, and the expression was outside the expected range of variation (|Z| > 4). As expected, empty vectors had no effect on the expression pattern and no HITs were found for this condition. A given gene was defined as HIT if at least one probe for this gene passed the threshold for a HIT. Based on random assignment of empty vector samples to groups of N=3 samples, we would expect 0-5 (mean 0.9) false positive gene HITS using our criteria.

### Gene-level effects and merging of platforms

Many probes are present for each gene on the Agilent array (one per exon for genes included) and a few on the Illumina array (one to three), with some of them performing better than the others and targeting different exons, and thus possibly different isoforms. For similarity analyses, a single value per gene was needed to represent the effect of ORFs. To compute these, the list of all probes flagged as HIT in any of the condition was extracted from the data, and the fold effect of all these probes for a given gene were averaged (geometric mean). The result is a matrix of effect sizes for all genes impacted by at least one ORF, computed for the whole list of ORFs. These values were used in downstream analyses such as computation of scores and similarity plots, to compare the effects of ORFs.

### Similarity Scores

Scores were computed as a projection of the effects of each ORF on the HITs of other targets ORFs. The score for ORF_1_ on ORF_0_ (the reference) is computed as the product of the (log) effects from both ORFs on the list of HITS from ORF_0_, scaled to set the score of ORF_0_ on itself to 1. The higher the score, the more ORF_1_ captures the effect of the reference ORF_0_. A negative score means an opposite effect. A score of 1 would be expected if ORF_1_ entirely capture the effect of ORF_0_. The scores are not symmetric, as they are computed using only the list of hits from the reference ORF. Only ORFs with at least 20 HITs were considered as reference ORF. This threshold of 20 was chosen to ensure sufficient power for interpretation of in-silico analyses and correspond to the (rounded) median the number of hits, after which we observe a steep increase in the distribution. At this threshold, we also expect that 95% of hits are true positives, thus reducing noise in in-silico analyses.

### Similarity plots

A two-dimensional projection of the similarity in effects between the ORFs was generated using the t-SNE method (t-Distributed Stochastic Neighbor Embedding) as implemented in R (library Rtsne) (30–32). The method is a non-linear data reduction algorithm aiming at finding non-linear structures and clusters in high-dimensional data. The projection was made from the gene-level effects matrix on the log scale, including ORFs with a least 20 HIT genes and with perplexity parameter set at 10. Given the random nature of the method, 30 independent projections were averaged and then used as the starting point of a last run of the t-SNE algorithm. The plot was generated using the igraph package (33), using the positions from the t-SNE projection of similarity in effects. Arrows were added to represent the pairwise similarity score.

### Bioinformatic annotation of HITS identified in the screen

In terms of cis-regulatory motif analyses of the proximal promoters of HITS identified in the screen, we used the PRIMA method (34) as implemented in the EXPANDER software (v8.0) (35). Specifically, we performed enrichment analyses aimed at detecting cis-regulatory DNA motifs (Jasper database) that were over-represented in the promoter sequences (Ensembl database v89) of the HITS for each of the ORFs expressed in HT-29; only enrichment of greater than 2-fold with a corrected P-value<0.001 (using the false discovery rate method) were considered. In order to find biological annotation categories enriched in HITS identified in the screen, we performed enrichment analyses using the g:GOSt functional profiling tool (using Ensembl release 96) from the online g:Profiler service (https://biit.cs.ut.ee/gprofiler/gost) (36, 37). Specifically, we evaluated enrichment for Gene Ontology (GO) terms (Biological process (BP), molecular function (MF) and cellular compartment (CC); release 2019-06-01) and biological pathways (KEGG, release 2019-06-03; Reactome, release 2019-06-05; and WikiPathways, release 2019-05-10); only enrichment with a corrected P-value<0.05 (using the g:SCS algorithm intrinsic to gProfiler which takes into account hierarchically related terms) were considered.

### qPCR validation of the effect of IFIH1

For the validation of the effect of IFIH1 in HT-29 cells and of the impact of the IBD-associated non-synonymous IFIH1 coding variants (see main text), RNA samples (n=3) from the microarray-based transcriptomic evaluation of IFIH1 and its three mutants were used to synthesize cDNA (as described above). The cDNA was amplified by qPCR using the PowerUp SYBR Green Master mix reagent according to the manufacturer’s recommendations (ThermoFisher Scientific) and with the QuantStudio 6 thermal Real time PCR system using the following cycle program: an incubation of 2 minutes for uracil-DNA glycosylase (UDG) activation at 55 ° C, an incubation of 2 minutes at 95 ° C for denaturation, followed by 40 cycles of 15 seconds at 95 ° C for denaturation, 30 seconds at 55 ° C for annealing and 60 seconds at 72 ° C for elongation, then a final cycle of 15 seconds at 95 ° C for denaturation, 60 seconds at 55 ° C for annealing and 15 seconds at 95 ° C for dissociation. The induction of the top 10 HITS from IFIH1 were validated in these samples via qPCR (PCR primer sequences used are shown in **Additional file 1: Table S3;** primers were obtained from Millipore Sigma unless mentioned otherwise from Integrated DNA Technologies). Relative expression data were normalized to expression of the Hypoxanthine Phosphoribosyltransferase (HPRT) gene.

### qPCR validation of the effect DUSP16

For the validation of the effect of DUSP16 and KSR1 in HT-29 cells, independent transductions were performed (as described above) in triplicate, and stable cell lines were established following puromycin selection; from these, total RNA was extracted, and cDNA synthesized (as described above). The induction of the top 10 HITS from DUSP16 and KSR1 were validated in these samples via qPCR (PCR primer sequences used are shown in **Additional file 1: Table S3** primers were obtained from Millipore Sigma unless mentioned otherwise from Integrated DNA Technologies). Relative expression data were normalized to expression of the HPRT gene.

### Functional analysis of DUSP16

For the validation of the functional impact of DUSP16 in HT-29 cells, a DUSP16 shRNA knock-down model and a Doxycycline-inducible model for DUSP16 expression in HT-29 cells were generated. Briefly, for the Doxycycline-inducible model, the lentiviral Lenti-X™ Tet-On® 3G Inducible Expression System (Takara Bio) was used. The codon-optimized DUSP16 ORF sequence was cloned into the lentiviral Lenti-X™ Tet-On® 3G Inducible response vector (Takara Bio) using the GATEWAY® cloning system and a lentiviral stock of this inducible DUSP16 expression plasmid was generated as described above. Following this, HT-29 cells were transduced with the pLVX-Tet3G regulator vector (Takara Bio USA) and a stable Tet-On 3G transactivator-expressing clonal HT-29 cell line (HT-29-pLVX-Tet3G) was isolated following selection with 500 ug/ml Geneticin (cat no 10131-027, Thermofisher Scientific). Lentiviral stocks for the pLVX-Tet3G regulator vector were produced as described above. The HT-29-pLVX-Tet3G line was then transduced in triplicate with either the lentiviral Lenti-X™ Tet-On® 3G Inducible response vector containing the DUSP16 ORF or the equivalent empty Lenti-X™ Tet-On® 3G Inducible response vector, and stable cell lines were obtained following selection with 400 ug/ml zeocin (cat no 46-0509, Thermofisher). The resulting Doxycyclin-inducible HT-29 cell lines were maintained in culture at 37°C with 5% atmospheric CO_2_ in McCoy’s 5A (Modified) medium (cat no 317-010-CL, Wisent Bioproducts) supplemented with 10% FBS premium quality, tetracycline free (cat no 081150, Wisent Bioproducts), 1X Glutamax (cat no 35050-61, ThermoFisher Scientific), 500 ug/ml Geneticin (cat no sc-29065A, Santa cruz Biotechnology), 100 ug/ml zeocin (cat no 46-0509, ThermoFisher Scientific). All HT-29 lines were maintained in the exponential growth phase (50-60% confluence) and used at 70-90% confluence for specific treatments and stimulations.

For the HT-29 shRNA knockdown model, five independent DUSP16-targetting shRNA lentiviral vectors from the MISSION® library (Sigma) were transduced in HT-29 cells. Lentiviral stocks for DUSP16 shRNA-expression vectors were produced following the same procedure described above. Stable shRNA DUSP16 knockdown cell lines were derived in triplicate for each shRNA vector following selection with 3 ug/ml puromycin and were maintained at 37°C with 5% atmospheric CO_2_ in McCoy’s 5A (Modified) medium (cat no 317-010-CL, Wisent Bioproducts) supplemented with 10% FBS (cat F1051-500mL, Wisent Bioproducts), 3 ug/ml puromycin (cat no P9620-10 ml, Millipore Sigma), 100 U/ml of penicillin and streptomycin (cat no 450-201-EL, Wisent Bioproducts) and 1X Glutamax (cat no 35050-61, ThermoFisher Scientific). Total RNA was extracted, and cDNA synthesized as described above. DUSP16 levels were evaluated by qPCR and the shRNA clone with the strongest impact on endogenous DUSP16 expression levels (TRCN0000052017) was selected to use in further experiments.

Specific conditions used in functional assays to induce DUSP16 ORF expression, as well as specific treatments (MAPK inhibitors and TNF-α stimulation) of the shRNA knockdown and DUSP16 expression models are described in the figure legends.

### Evaluation of the effect of DUSP16 expression in HCT-15 cells

Proliferative HCT-15 cells at 50% confluency (in 12 well plate (Corning-Thermo Fisher Scientific)) were transduced in triplicate with viral stock of pLVX-EF1a-IRES-PURO/eGFP, either empty or containing the DUSP16 ORF (as described above) and stable cell lines were established following puromycin selection. Specific conditions and treatments used in functional assays (MAPK inhibitors and TNF-α stimulation) of this DUSP16 expression HCT-15 model are described in the figure legends.

### Statistical analyses for functional analyses

Expression obtained by qPCR was normalized by the endogenous gene HPRT and log2-transformed. When different plates were used, systematic plate effects were removed by the mean across all conditions (or median if there was large variation between conditions). When many treatments were tested on the same cell lines, systematic cell line differences were removed by the mean across all treatment. The mean and standard error of the mean (SEM) for each condition were computed and transformed back to the original scale as geometric mean (estimator of population median) with SEM, with all data points shown. In the context of increased concerns on the overuse (and misuse) of P-values in reporting biological data, and given the nature of this project, we decided to use inference on parameters instead of reporting p-values from statistical testing of the null hypothesis. As an indication, non-overlapping three datapoints from two conditions correspond to the strongest evidence against the hypothesis of no difference for a non-parametric test (P-value = 0.1). Under hypothesis of lognormal distribution, the intervals as plotted correspond to 58% confidence interval for the median.

## RESULTS

### Lentiviral ORF screen in human colonic epithelial cells

To better understand the functional impacts of IBD genes in disease susceptibility within the intestinal epithelial cell compartment, we performed an expression-based screen of genes within known IBD loci (see overview of study flow in **Additional file 2: Fig.S1)**. In order to define the gene list to include in this transcriptomic screen, we began with the 167 IBD-associated loci identified, just prior to the initiation of the current project, by the International IBD Genetic Consortium’s genetic analysis of ∼75,000 IBD cases and controls (1). The 297 genes found within these loci were evaluated for their expression profile in a panel of human tissues and cell lines. With this information we prioritized a list of 169 genes from IBD-associated loci with intestinal expression, including 20 known causal genes, for our expression-based screen in human colonic epithelial cells (**Additional file 1: Table S1**). The next step in the experimental design was to clone an open reading frame (ORF) for each of these genes into a lentiviral expression vector, and then transduce each into HT-29 cells via three independent infections. Over 85% (147/169) of these ORFs were successfully cloned, transduced, and generated stable cultures. Following an average of 8 days in culture to select successfully transduced cells and obtain confluent cultures, RNA was extracted, and the transcriptomes of the different stable cultures were assessed on two different microarray expression profiling platforms (**Additional file 2: Fig.S1)**.

After normalisation of this transcriptomic data, merging data from samples of different experimental batches and merging of experimental replicates (independently on each of the microarray platforms), we first assessed the expression level of each transduced ORF in the cell’s transcriptome (**Additional file 1: Table S4 & Additional file 2: Fig.S2**). We then excluded two ORFs that did not show detectably increased expression and set out to determine the impact of the remaining 145 ORFs on the cell’s transcriptome. To do so, we determined the variance in expression of all detectable genes in the transcriptome across all 435 samples included in these analyses, and then determined the set of genes that increased or decreased in response to each ORF– these “HITS” were defined as genes where the fold effect on their expression levels (either increased or decreased) computed from their combined replicates over their median expression was larger than two-fold and their expression was outside their expected range of variation (**Fig.1A**). Results from the two expression profiling platforms were combined (see Methods) before proceeding to in silico analyses (**Additional file 2: Fig.S1**).

**Fig.1:**
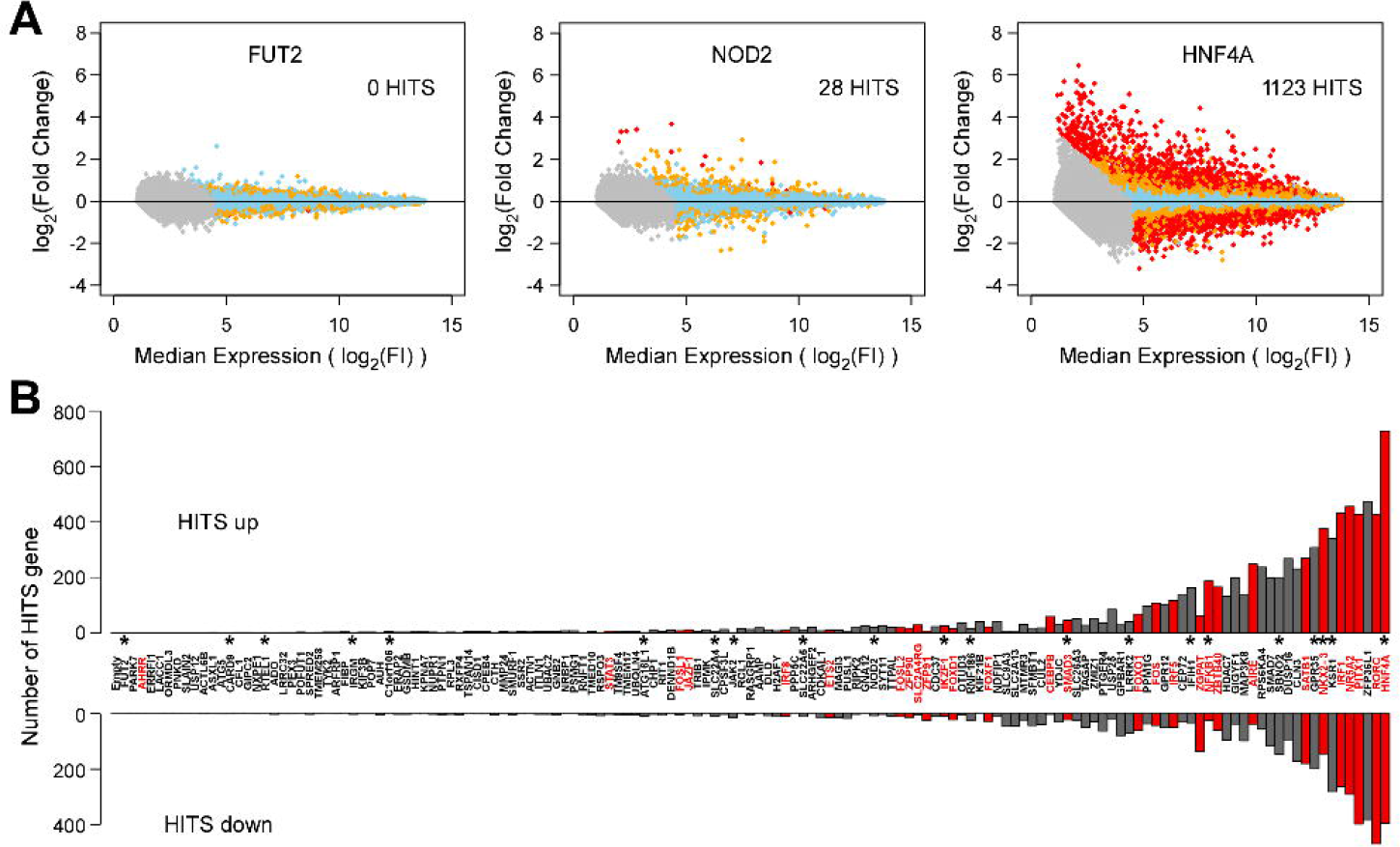
Impact of IBD gene candidate ORFs on the HT-29 transcriptome. **(A)** Selected examples illustrating the impacts observed on the transcriptome of HT-29 cells following the expression of different ORFs for IBD gene candidates. HITS are identified as genes with probes from either microarray platform showing detectable expression in HT-29 (endogenously or following ORF expression) for which the fold effect in response to the expression of a given ORF is greater than two compared to the baseline, and shows expression outside the expected range. As examples, FUT2 (left), a terminal enzyme in a metabolic pathway, had no impact on the transcriptome (0 HITS); NOD2 (middle), an intracellular PAMP receptor, had only marginal impact on the transcriptome (28 HITS) in the absence of its ligand; and HNF4A, a transcription factor known to have a central role in intestinal epithelial cells, had the strongest effect (1123 HITS). Each dot represents a single detectable probe from the genome-wide array tagging a specific gene in the HT-29 transcriptome. The x-axis shows the log_2_-transformed median expression across all conditions (baseline). The y-axis represents the effect of transduction of a given ORF, as the log_2_-transformed fold-induction compared to baseline. Skyblue dots are probes with expression value within expected range of variation ( |Z| ≤ 2), orange dots represent probes suggestively outside the range ( |Z|>2 ) and red dots represent probes outside the range ( |Z| > 4). Gray dots are probes with expression value below detection threshold. **(B)** Impact of the expression of all IBD gene candidates on the transcriptome of HT-29 cells. ORFs are ordered along the x-axis by increasing total number of HITs (see **Additional file 1: Table S5**), with number of up- and down-regulated HITs gene shown along the y-axis. Starred ORFs are previously reported IBD candidate causal genes and ORFs listed in red indicate known transcription factors (as defined by **Lambert et al.** (**90**)).

Importantly, low variances were observed between the replicates for any given ORF, highlighting the robustness of the experimental approach used (median SD between log_2_(FC) was 0.12, with an interquartile range of 0.35). As could be expected, the empty vector had no effect on the expression pattern and no HITS were found for this condition. Moreover, the different ORFs showed a wide range of effects on the cell’s transcriptome (***from 0 to 1123 HITS***), with ORFs encoding for transcription factors or for molecules involved in intracellular signalling (e.g. G protein-coupled receptors, phosphatases and kinases) showing the greatest impact on the transcriptome (**Fig.1B** **& Additional file 1: Table S5**. ORFs encoding structural proteins, terminal enzymes in metabolic pathways or proteins whose function likely requires an external stimulus, such as for the known causal genes C1ORF106, FUT2 and IRGM, respectively, had little impact on the transcriptome.

As a first-pass validation of the observed impact on the transcriptome, a subset of 9 ORFs were repeated, each with an additional three independent infections performed later in time (4 to 16 months). Results were highly consistent between dates (**Additional file 2: Fig.S3**). Taken together, these analyses suggest that this approach is robust, and shows reproducible ORF-dependent impacts on the HT-29 transcriptome.

### HNF4A has a major impact on the epithelial transcriptome

As a first step to obtaining a biological interpretation of the HITS emanating from these analyses, we examined the results of HNF4a as it is the ORF showing the strongest effect in our epithelial model with 1176 HITS (728 genes that increased and 395 genes that decreased in their expression). HNF4a is a known transcription factor previously described to have a prominent role in the differentiation and function of hepatic, pancreatic and intestinal epithelial cells (38–40), and is believed to be the causal gene within its susceptibility locus (41). In our experiment, the transduction of the HNF4a ORF led to an increase of 13-fold of the HNF4a transcript in HT-29 cells. The top five upregulated genes were the ABCB1, MPP1, SI, SPRR3 and ANPEP genes, which are known to play important roles in intestinal transport of xenobiotics, cell junctions and polarity, digestion of dietary carbohydrates, epithelial cell structure, and digestion of peptides, respectively (42–46). A global annotation analysis of the 728 genes that increased following expression of HNF4a found significant enrichment of multiple annotation terms, most of which are known to play essential roles in intestinal epithelial function such as multiple metabolic processes, transmembrane absorption & transport of lipids, alcohols, organic acids, steroids, small molecules, ions, drugs & xenobiotics, in addition to the formation of specialized structures of the apical plasma membrane (microvilli and brush border) and the regulation/resolution of inflammatory responses (**Additional file 2: Fig.S4A-D**).

In addition, an analysis of the proximal promoters of these 728 genes revealed a significant enrichment for the HNF4a transcription factor binding site (TFBS) (**Additional file 1: Table S6**). We have previously reported a connection between HNF4a and RNF186, another causal IBD gene, specifically hypothesizing that HNF4a can regulate the expression of HNF1a, which then regulates the transcription of RNF186 (5). Indeed, in the current dataset HNF1a and RNF186 transcripts are increased by 2.4- and 5.4-fold, respectively, in HT-29 lines expressing the HNF4a ORF. Taken together, these results support that our experimental approach has the capacity to identify ORF-related functions relevant to epithelial biology.

### Identifying Shared, Distinct, and Opposing Impacts of ORFs on Epithelial Transcriptome

In order to systematically screen through our transcriptomic dataset of 145 IBD ORFs and identify genes that share an impact on similar biological functions or processes, we developed a “similarity score” approach to cluster genes with similar effects on the transcriptome. We focused our analysis on ORFs that had at least a total of 20 HITS that were either up- or down-regulated. This threshold of 20 was chosen to ensure sufficient power for interpretation of in-silico analyses and corresponds to the (rounded) median the number of HITS across all conditions, after which we observe a steep increase in the distribution. As can be seen in **Fig.2**, this analysis revealed two large distinct gene clusters: ***Cluster 1*** includes multiple known IBD causal genes including IFIH1, SBNO2, NFKB1 and NOD2, with several candidate genes connecting to them, while ***Cluster 2*** has multiple candidate genes connecting to the IBD causal gene KSR1.

**Fig.2:**
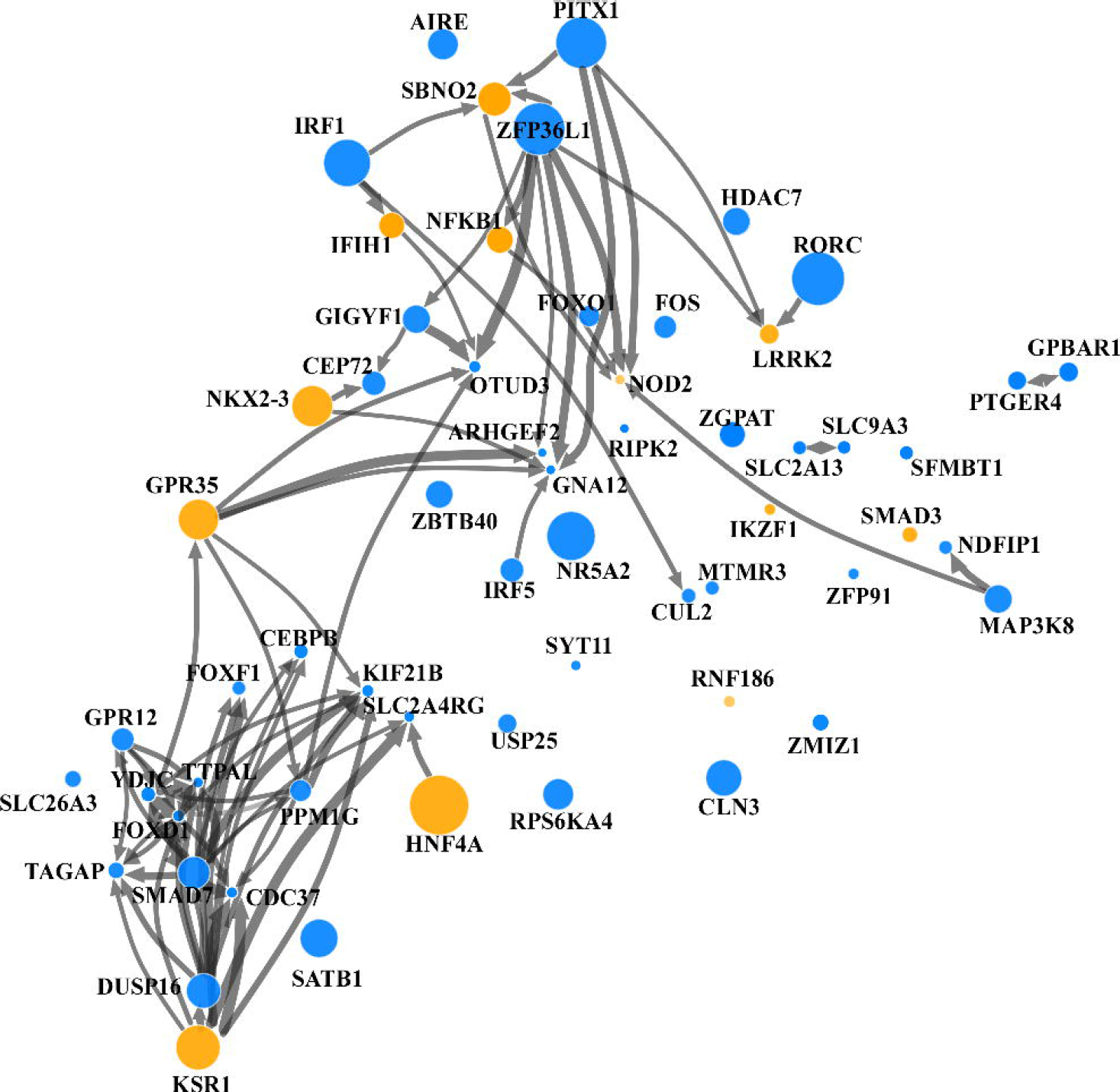
Clustering ORFs based on shared HITs and pairwise similarity of effect on the HT-29 transcriptome. A similarity analysis was applied to the transcriptomics dataset from our expression screen for IBD gene candidates in HT-29 cells. A two-dimensional projection of the similarity in effects between the ORFs was generated using the t-SNE method from the gene-level effects matrix on the log scale, including ORFs with a least 20 up-regulated or down-regulated HIT genes. ORFs with a similar effect on the transcriptome are clustered together on the graph. Size of the circle for each ORF was made proportional to the number of HITS. Yellow circles represent IBD-causal genes. Arrows were drawn between ORFs with width proportional to a similarity score (see methods). Arrows from ORF_1_ to ORF_0_ represent evidence of shared effects on the subset of the transcriptome defined by the HITS of ORF_0_. Only arrows for a positive score greater than a specific threshold (0.6) were drawn. Two clusters can be loosely defined in top right (cluster 1) and bottom left (cluster 2) with little connections between them

This analytical approach allows us to identify genes with shared impact on the transcriptome, as further illustrated for **Cluster 1** genes by pairwise comparisons of causal IBD genes IFIH1, SBNO2, NFKB1, and NOD2 showing highly correlated effects on the epithelial cell’s transcriptome (**Fig.3A**). Similarly, for **Cluster 2**, IBD-causal gene KSR1 shares a comparable impact on the transcriptome to IBD gene candidate DUSP16 and known causal gene HNF4A (**Fig.3B**).

**Fig.3:**
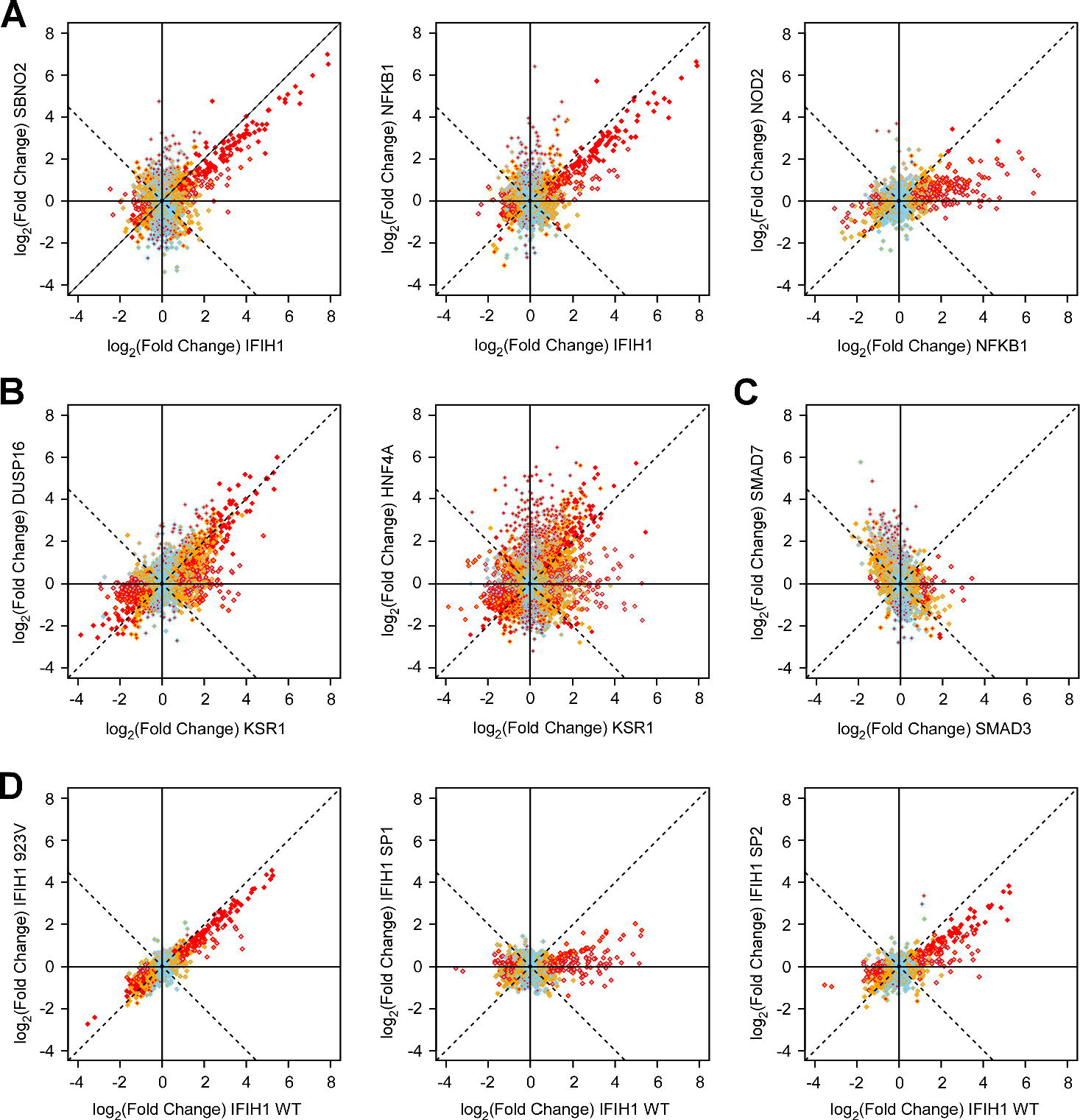
Correlation of effect of different IBD gene candidate ORFs on the HT-29 transcriptome. Correlation plots illustrating the shared effects of IBD gene candidate ORFs on the transcriptome of HT-29 cells are shown **(A)** for cluster 1 IBD causal genes IFIH1 vs NFKB1 and SBNO2, and NOD2 vs NFKB1, **(B)** for cluster 2 IBD causal gene KSR1 and its closest candidate ORF cluster neighbor DUSP16, as well as KSR1 causal gene HNF4A, and **(C)** for IBD causal gene SMAD3 and candidate ORF SMAD7. Additional correlation plots for IBD-causal genes with IBD gene candidate ORFs are shown in **Additional file 2: Fig.S6**. **(D)** Correlation plots comparing the effects of increased wild-type IFIH1 expression against expression of three of its IBD-associated non-synonymous coding variants: an isoleucine to valine substitution at position 923 (rs35667974) (923V, **left**), a splice donor site variant at position +1 in intron 8 (SP1, **middle**) and a splice donor site variant at position +1 in intron 14 (SP2, **right**). For the effect of IFIH1 variants on IFIH1 HITs, see **Additional file 1: Table S9.** For all correlation plots shown, each dot represents a single detectable probe from the genome-wide array tagging a specific gene in the HT-29 transcriptome (see **Fig.1**). The x-axis (inner color of dots) and y-axis (border color of dots) show the effect of two independent set of replicated ORFs on the transcriptome, as the log2-transformed fold-induction compared to baseline. Skyblue are probes with expression value within expected variation (|Z| ≤ 2), orange represent probes suggestively outside the range (|Z|>2 ), red represent probes outside expected range of variation (|Z| > 4), and g. Gray are probes with expression value below our detection threshold; this color code is applied to the inside of dots for x-axis data and the border of dots for y-axis data. No metric of correlation was included, as most probes are not affected by ORFS and thus only represent noise. See **Fig. 2** for similarity illustration.

This graphical representation can also highlight genes that have very distinct effects, with few to no connections with others. One example is the IBD-causal gene SMAD3, whose impact on the transcriptome is without any noticeable resemblance to others. SMAD3 is a key molecule in the signaling cascade of the TGF-β receptor. Interestingly, IBD-candidate gene SMAD7, which is very distant in the similarity graph from SMAD3, is a direct inhibitor of SMAD3 signaling and is known to block the effect of SMAD3 in response to IFN-γ (47). The opposite effects of the agonistic SMAD3 and antagonistic SMAD7 are further illustrated in a direct comparison of their impact on the HT-29 transcriptome (**Fig.3C**).

### Multiple ibd genes are involved in the immune response of epithelial cells to the local microbial environment

As described above, ***Cluster 1*** included four known causal IBD genes (IFIH1, SBNO2, NFKB1 and NOD2) with similar effects on the transcriptome. IFIH1 encodes MDA5, an innate pattern recognition receptor (PRP) that functions as a cytoplasmic receptor for viral RNA (48), which upon binding to RNA leads to the activation of IRF3/7 and the transcription of type I interferon genes (**Additional file 2: Box 1**) (49). The encoded interferon proteins can then act in an autocrine and/or paracrine fashion, triggering multiple signaling cascades (e.g. JAK-STAT, NFkB etc.) that result in the transcriptional regulation of hundreds of genes, many of which have interferon-sensitive response elements (ISRE), that bind the heterodimeric transcription factor (STAT1::STAT2), or NFkB binding sites in their promoters (50). Analysis of the promoters of the genes activated by the expression of these four IBD genes (**Additional file 1: Table S6)** found an enrichment of binding sites for STAT1::STAT2 and for members of the IRF family in three of these (IFIH1, SBNO2, and NFKB1) and of NFkB sites (REL and RELA) in two (NFKB1 and NOD2). These TFBS-based analyses support a role for IFIH1, SBNO2, and NFKB1 in the anti-viral and type 1 interferon responses, and for NFKB1 and NOD2 a role in anti-bacterial responses (**Additional file 2: Box 1**). The enrichment of TFBS increases as a function of the stringency of threshold for defining a HIT from the screen (**Additional file 1: Table S6**). These observations are further supported by enrichment analyses of functional annotations in the HITS for SBNO2, NFKB1, and IFIH1(**Additional file 3: Cluster 1 data**), which showed a very important shared enrichment for anti-viral responses as well as responses to type I interferons (**Additional file 1: Table S7)**, and significant shared enrichment of chemokine-driven responses in the HITS of NFKB1 and NOD2 (**Additional file 1: Table S8**). These results are entirely consistent with the known functions of IFIH1, NFKB1, and NOD2 (**Additional file 2: Box 1**) (48, 51, 52). While SBNO2 is considered a causal IBD gene, its function is not well characterized although first identified as a component of the IL-10 signaling cascade in monocytes (**Additional file 2: Box 1**) (53).

As an additional validation of our experimental approach’s ability to capture biologically relevant effects, we were interested in examining the impact of IBD-associated variants in the known IBD causal gene IFIH1 in our HT-29 model. The IFIH1 region was first identified as an IBD risk locus in a meta-analysis of GWAS, and subsequent high-density SNP based association mapping identified a single non-synonymous variant in this gene (rs35667974, I923V) associated to UC risk with >95% certainty, confirming this as the most likely causal gene in the region (8). In a previously published targeted sequencing of the exons of 759 protein-coding genes from IBD loci, we found two additional previously reported coding variants in the IFIH1 gene associated with increased risk to IBD: rs35337543, at splice donor site position +1 in intron 8 leading to a predicted skipping of exon 8 and an in-frame deletion within the first helicase domain (MAF=1%, OR=1.3, P-value=0.003), and rs35732034 (MAF=0.8%, OR=1.3, P-value=0.002), at splice donor site position +1 in intron 14 leading to a predicted skipping of exon 14 and a premature termination resulting in the truncation of the C-terminal domain (CTD), which recognizes and binds viral RNA (54, 55). We therefore cloned the resultant variant ORFs and expressed these in HT-29. As can be seen in **Fig.3D** **and Additional file 2: Fig.S5A**, these IBD-associated variants had an impact on the transcriptome that was quantitatively (**Additional file 1: Table S9)**, but not qualitatively different as compared to the wildtype ORF, either from a global perspective (**Fig.3D**) or when evaluating the top 10 HITS in the cells expressing the IFIH1 ORF (**Additional file 2: Fig.S5A)**. The microarray results for these same top 10 IFIH1 HITS were also further validated by qPCR (**Additional file 2: Fig.S5B)**.

Extending our analyses to the remaining genes within **Cluster 1**, also identified highly-significant enrichments of IRF1, IRF7, and STAT1::STAT2 sites within the promoters of the HITS from ZFP36L1, IRF1, GIGYF1, OTUD3, and AIRE expressing lines, and of NFkB sites within the promoters of the HITS from PITX1 expressing lines, but no detectable enrichment of such sites for FOS or FOXO1 HITS (**Additional file 1: Table S6**). This pattern was also seen in the enrichment of functional annotations (**Additional file 3: Cluster 1 data**). Specifically, the HITS of ZFP36L1, IRF1, GIGYF1, OTUD3, and AIRE were enriched for shared functional annotations related to anti-viral responses and Type-1 interferon response (**Additional file 1: Table S7)**, the PITX1 HITS being enriched for shared cytokine- and chemokine-driven responses (**Additional file 1: Table S8)**, and the HITS of FOS and FOXO1 being more modestly enriched for a broad range of functional annotations. These shared functions are in part due to shared HITS, as illustrated by correlation plots between the different cluster 1 IBD-causal genes and IBD gene candidate ORFs (**Additional file 2: Fig.S6A)**, for example over a third of NFKB1’s HITS (67 of 185) are in common with PITX1.

We therefore propose that multiple IBD genes play an important role within the intestinal epithelium in terms of transcriptional regulation of the type I interferon response to viral pathogens (SBNO2, NFKB1, IFIH1, ZFP36L1, IRF1, GIGYF1, OTUD3, and AIRE) and others in the transcriptional regulation of the anti-bacterial response (NFKB1, NOD2, PITX1). While for many of these genes (e.g. IFIH1, NFKB1, IRF1, NOD2, FOS) the association with anti-microbial functions has been well established, for the others this association is novel (**Additional file 2: Box 1**).

### KSR1 and DUSP16 play a role in intestinal homeostasis

**Cluster 2** contains ten genes from ten different IBD loci, with only KSR1 having previously been established as causal (**Additional file 1: Box 2**). KSR1 encodes for a protein that has active kinase activity and that acts as a scaffold for the Ras/Raf/MAPK signaling pathway (56). While KSR1 is most often studied within the context of oncogenesis, GWAS studies of IBD and subsequent high-density SNP based association mapping implicated KSR1 as a causal IBD gene (1, 8). Moreover, multiple studies have linked KSR1 to inflammation (**Additional file 2: Box 2**). In the current screen, KSR1 had 340 HITS and consistent with its broad role in signaling, functional annotation of these HITS found enrichment for a large number of biological processes and functions such as “metabolism of fatty acids”, “biosynthesis of specialized pro-resolving mediators”, “regulation of gene expression in endocrine-committed (NEUROG3+) progenitor cells”, and “drug metabolism”, all known functions of intestinal epithelial cells (**Additional file 5: Cluster 2 data**). In fact, 50% of the annotation terms enriched in KSR1’s HITS are shared with HNF4A, including metabolism of fatty acids, biosynthesis of specialized pro-resolving mediators, formation of brush border membrane and microvilli, and drug metabolism, which is further supported by the fact that KSR1 shares 102 (30%) of its 340 HITS with the well-characterized HNF4A (**Fig.3B**). This suggests that KSR1 plays an important role in regulating intestinal epithelial function.

Of the 10 genes contained in cluster 2, CDC37, DUSP16, TAGAP, SMAD7 and SATB1 share respectively 65%, 44%, 40%, 27% and 21% of their HITS with KSR1 (**Additional file 2: Fig.S6B)** suggesting that these all share some role in intestinal epithelial function. DUSP16, which was the closest to KSR1 in the similarity network (**Fig.2**) and in a pairwise comparison (**Fig.3B**), had 119 of its 268 HITS in common with KSR1 and shared many functional annotations terms such as “organic acid metabolism”, “brush border cellular compartment”, and “regulation of gene expression in endocrine-committed (NEUROG3+) progenitor cells (**Additional file 1: Table S10)**. It is important to note that while genes may share functional annotations, the actual HITS may be different. For example, while DUSP16 and KSR1 both show enrichment for the GO annotation term for proteins involved in the “brush border” cellular compartment, and the HITS for KSR1 and DUSP16 both include eight genes tagging this GO annotation term, only four of these are in common (SI, CDHR5, CYBRD1, PDZK1). Interestingly, this annotation term is also highly enriched in the HITS from HNF4A expressing lines, suggesting that KSR1 and DUSP16 share a role with HNF4A in regulating some key homeostatic functions of intestinal epithelial cells, although not always impacting on the same list of genes.

The observation that KSR1 and DUSP16 have shared functions is further supported by the overlap of six of the top ten HITS of KSR1 (in order <u>CHP2</u>, <u>C15orf48</u>, <u>FCGBP</u>, <u>ENPP2</u>, <u>SI</u>, AKR1C2, CYP1A1, <u>HEPACAM2</u>, NR1H4, RETREG1) and of DUSP16 (in order <u>CHP2</u>, <u>C15orf48</u>, <u>ENPP2</u>, <u>SI</u>, CASP1, PIGR, FCGBP, SST, <u>HEPACAM2</u>, MS4A8). Moreover, even for the top 10 genes that are not shared between KSR1 and DUSP16, most can be found within the broader set of HITS for these genes. Importantly, these top 10 HITS alone highlight some important intestinal epithelial functions such as dietary metabolism (SI, NR1H4), inflammatory processes (CASP1, SST), as well as homeostasis and defense (CHP2, PIGR, FCGBP, HEPACAM2); see **Additional file 1: Table S11** for details. As outlined in **Additional file 1: Table S11**, several of these genes are also found within the HITS of other Cluster 2 genes.

In order to validate these observations, we independently repeated the transduction of KSR1 and DUSP16 in HT-29 cells and validated the impact of these ORFs on their top HITS (**Additional file 1: Table S12**). Given that DUSP16 functions as a dual specificity protein phosphatase involved in the inactivation of MAPK (57, 58), it is reasonable to think that DUSP16 is impacting the transcriptome of epithelial cells via regulation of MAPK activity. To test this hypothesis, we chose to focus on the potential link between DUSP16 and the PIGR gene that is suggested by our transcriptomic data, as it encodes the pIgR receptor that is known to have a role in the protection of the mucosa from intestinal microbiota and is impacted by several other IBD gene candidates (KSR1, SATB1, IRF1, and GIGYF1)(**Additional file 1: Table S11**).

### The novel IBD gene DUSP16 functions as a regulator of MAPK activity and is linked to epithelial protectioN

We first generated stable DUSP16 shRNA knock-down HT-29 cell lines (with 80% KD efficiency on RNA) (**Additional file 2: Fig.S7A**) and confirmed that reduction of DUSP16 levels leads to a 60% reduction of basal pIgR expression levels (**Additional file 2: Fig.S7D**). In order to examine the kinetics of the effect of DUSP16 on PIGR expression in a dose- and time-controlled manner, we generated a tetracycline-inducible model for DUSP16 expression in HT-29 cells co-expressing the Tet-On 3G transactivator. Using this model, we were able to obtain controlled DUSP16 expression levels (from 2- to 300-fold increase in expression over endogenous levels) over a wide range of doxycycline (DOX) concentrations (1-1000ng/ml) following a 24-hour treatment (**Additional file 2: Fig.S7B**). Interestingly, using the same samples we observed that PIGR expression levels were not linearly correlated to DUSP16 expression levels within the range of DOX concentration tested but rather seemed to sharply increase four- to five-fold when DUSP16 increased 60-fold (at a DOX dose of 10 ng/ml) and remained stable even at higher doses of DUSP16, indicating that the effect of DUSP16 on PIGR expression rapidly becomes saturated in these cells (**Additional file 2: Fig.S7E**). Using the lowest dose tested that induced maximal PIGR increase in our model (DOX dose of 10 ng/ml), we were able to show that DUSP16 achieved a steady state expression level within 24 hrs of DOX stimulation (**Additional file 2: Fig.S7C**) while PIGR continuously increased over time (from 1 to 12 days), reaching induction levels close to 300-fold (**Additional file 2: Fig.S7F**). The timing of PIGR induction closely mirrors the increase in levels of DUSP16, suggesting that the impact of DUSP16 on PIGR expression likely does not involve the induction of multiple of intermediary factors.

Based on these results and on the role of DUSP16 in MAPK regulation, we then tested the effect of DUSP16 expression on MAPK activity in our cell model. More specifically, we evaluated phosphorylation levels of ERK, p38 and JNK following induction of DUSP16 for 24 hours (**Fig.4A**). Previous reports have shown that DUSP16 can impact on MAPK with a higher specificity for JNK and p38 over ERK. As expected, expression of DUSP16 in proliferative cells strongly reduced steady-state phosphorylation levels of p38 and JNK but had a more marginal effect on ERK. In addition, we tested the impact of DUSP16 in a serum deprivation model followed by a stimulation of MAPK activity with TNF-alpha (**Fig.4B**). In this model, not only do we observe that DUSP16 can further reduce phosphorylation of all three MAPKs following serum deprivation, it can also inhibit activation of all three MAPKs following TNF-alpha stimulation. These results indicate that under the right conditions DUSP16 can not only impact on p38 and JNK activity, but on ERK activity as well.

**Fig.4:**
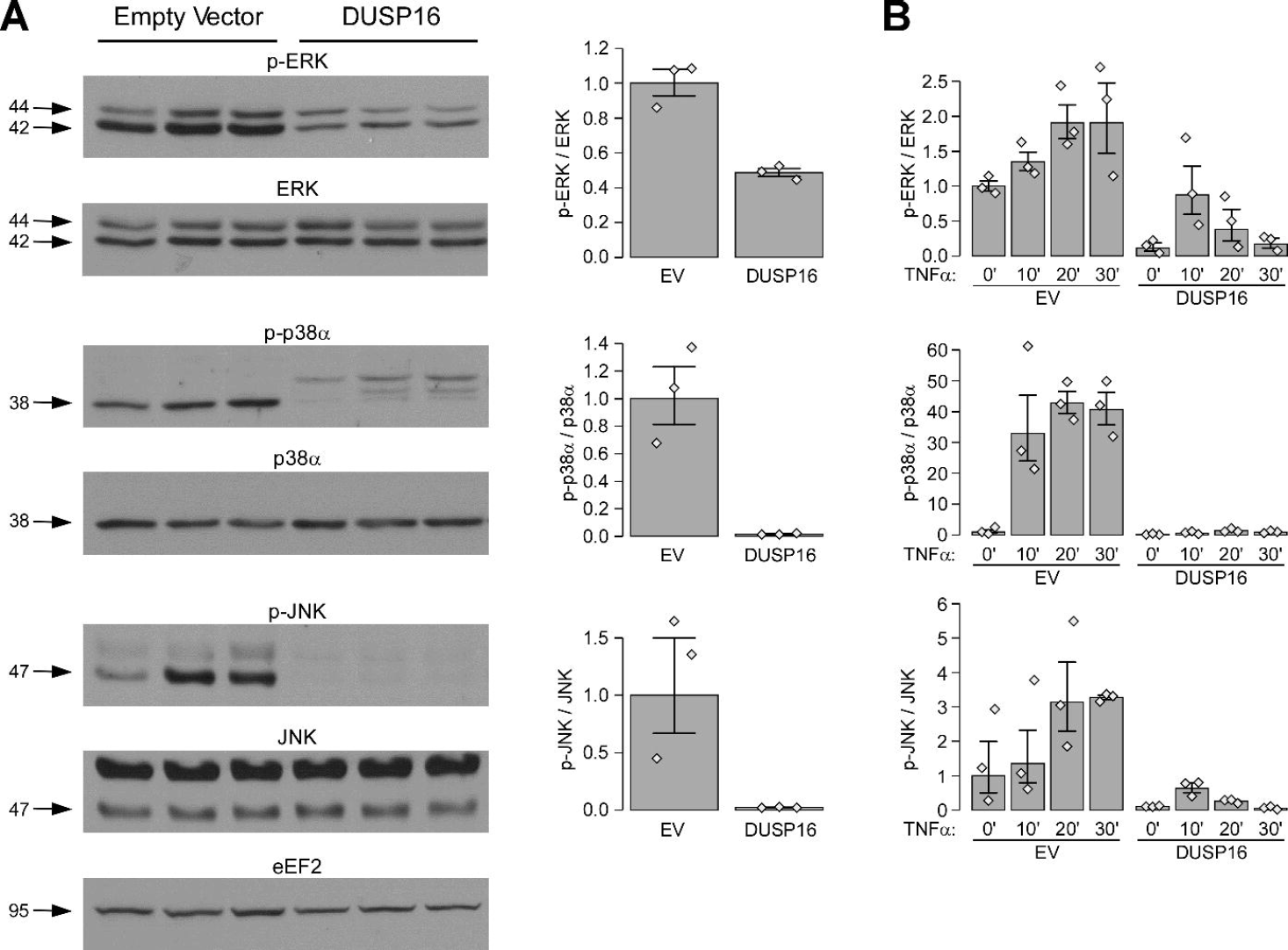
Impact of increased DUSP16 expression on MAPKs phosphorylation levels. The impact of DUSP16 expression on ERK, p38 and JNK phosphorylation levels was evaluated using a Doxycycline inducible HT-29 cell model. **(A)** The impact of DUSP16 on steady-state MAPKs phosphorylation levels was measured. Exponentially growing HT-29-pLVX-Tet3G cells stably transduced with a TET3G-inducible expression plasmid (empty vector (EV) or containing the DUSP16 ORF (DUSP16) were stimulated with Doxycycline (1ug/ml) for 24 hours and cell lysates were harvested for the evaluation of the native and phosphorylated states of MAPKs. Cropped western blots showing bands of interest are shown (left) and the level of phosphorylation of each MAPK is summarized in a graphical representation (right). A single representative gel is shown for EEF2 loading control; Full blots along with their respective EEF2 are shown in **Additional file 2, Fig.S11, panel A**. **(B)** The impact of DUSP16 on the induction of MAPKs phosphorylation following serum starvation was measured. Using the same models as above, exponentially growing cells were first serum-starved for 24 hours (0% FBS) in the presence of the Doxycycline inducer (1ug/ml). The cells were then stimulated with 10 ng/ml TNF-α for different times to induce MAPK phosphorylation and whole cell lysates harvested for the evaluation of the native and phosphorylated states of MAPKs. The level of phosphorylation of each MAPK is summarized in a graphical representation; the y-axis shows phosphorylation levels relative to the control condition (EV in **panel A** and EV 0’ in **panel B**). All phosphorylation level data is expressed as the ratio of phosphorylated over total MAPK (both corrected for EEF2 loading control) combining the replicates (n=3). All results are shown as geometric means with SEM. The values for independent replicates are shown (grey lozenges). As an indication, three non-overlapping datapoints from two conditions correspond to the strongest evidence against the hypothesis of no difference for a non-parametric test (P-value = 0.1). Under hypothesis of lognormal distribution, the intervals as plotted correspond to 58% confidence interval for the median. Full blots for each replicate and all timepoints, along with EEF2 results are shown in **Additional file 2, Fig.S11, panel B-C**.

In order to further characterize the mechanism by which DUSP16 impacts on PIGR regulation, we evaluated the effect of chemically inhibiting each MAPK kinase independently (**Fig.5A**). Using this model, we first observed that a short term (24 hr) chemical inhibition of ERK causes a marginal but detectable induction of PIGR similar to the one obtained via expression of the DUSP16 ORF (**Fig.5B**) with a similar effect also seen for chemical inhibition of p38 but not JNK. Moreover, a long-term chemical inhibition of ERK and p38, but not JNK, showed a very strong induction of PIGR expression (25- and 15-fold respectively). The effects observed from the chemical inhibition of MAPKs were similar to the effects seen following expression of the DUSP16 ORF for an equivalent period of time. Taken together, our results show that DUSP16 can repress p38 and JNK activity, while showing a marginal impact on ERK, and also show that inhibiting ERK and p38 can strongly induce PIGR expression, suggesting that DUSP16 leads to increased PIGR expression most likely predominantly through its inhibition of p38 but also more mildly through the modulation of ERK activity.

**Fig.5:**
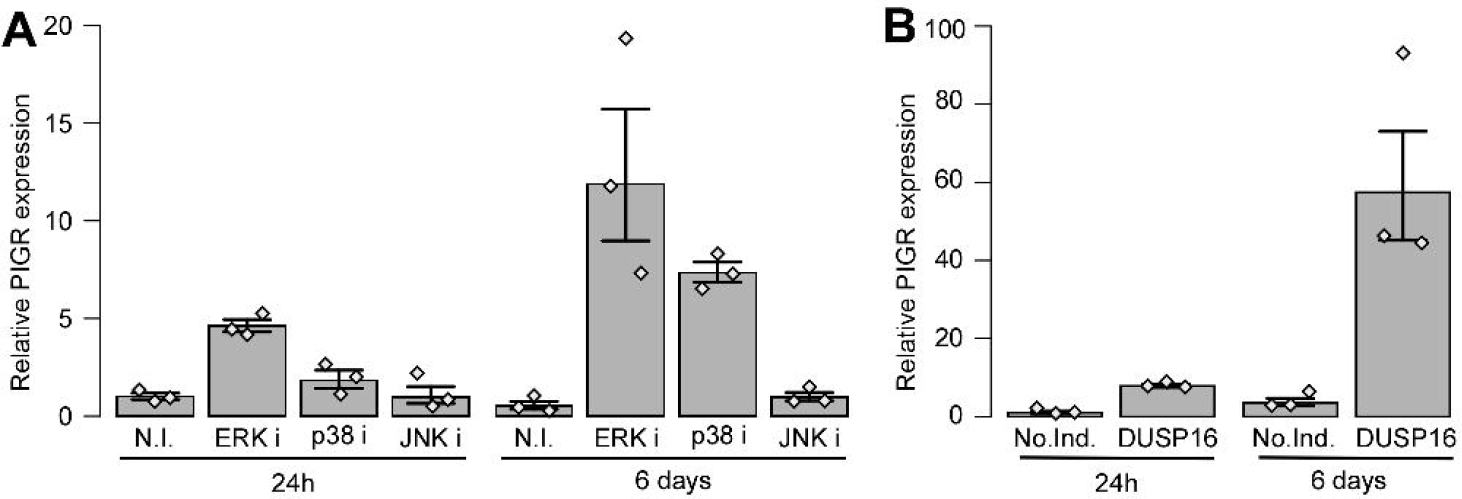
Impact of short- and long-term inhibitions of MAPKs on PIGR expression. The impact of short-term (24 hours) and long-term (6 days) inhibitions of MAPKs on PIGR mRNA levels was evaluated by qPCR. **(A)** Exponentially growing HT-29-pLVX-Tet3G cell lines (n=3) were either left untreated (no inhibitors; N.I.) or treated with chemical inhibitors specific for the different MAPKs (PD98059, ERKi; SB203580, p38i; SP600125, JNKi) at a concentration of 10uM, and total RNA was isolated at two different timepoints (24hours and 6 days). Expression levels of PIGR were evaluated via qPCR; the qPCR results from the replicates were combined and mean expression values relative to samples without inhibitors (N.I.) are shown. **(B)** Exponentially growing HT-29-pLVX-Tet3G cell lines stably transduced with the TET3G-inducible expression plasmid for DUSP16 ORF (n=3) were either left untreated (No Doxycycline induction; No.Ind.) or stimulated with Doxycycline at a concentration of 10 ng/ml to induce DUSP16. Cells were harvested at two different timepoints (24hours and 6 days) for total RNA isolation. Expression levels of PIGR were evaluated via qPCR; the qPCR results from the replicates were combined and mean expression values relative to samples without treatment with doxycycline (No.Ind.) are shown. All results are shown as geometric means with SEM. The values for independent replicates are shown (grey lozenges). As an indication, non-overlapping 3 datapoints from two conditions correspond to the strongest evidence against the hypothesis of no difference for a non-parametric test (P-value = 0.1). Under hypothesis of lognormal distribution, the intervals as plotted correspond to 58% confidence interval for the median.

A previously published report had shown that in a HT-29 subclonal line, selected for increased basal levels of PIGR (59), chemically inhibiting ERK and p38 had minimal impact on basal levels of PIGR; however, combining these inhibitions with a TNF-α stimulation caused a synergistic induction of PIGR expression (60, 61), that was not seen under inhibition of JNK. Based on this we set out to confirm these observations in our model and evaluate whether DUSP16 can have a similar effect to chemical MAPK inhibitors in our model. We confirm that in our model, combining TNF-α stimulation to ERK or p38 chemical inhibition leads to a synergistic induction of PIGR gene expression (**Additional file 2: Fig.S8A**); furthermore, we demonstrate that pIgR expression is also further increased in HT-29 cells expressing the DUSP16 ORF as compared to parental HT-29 cells in response to TNF-α stimulation, in the absence of chemical MAPK inhibitors (**Additional file 2: Fig.S8B**). Moreover, we show that PIGR induction by TNF-α is also abrogated when tested in our shRNA knock-down model where the endogenous levels of DUSP16 were reduced by 80% (**Additional file 2: Fig.S8C**). Taken together, our data suggests that DUSP16, through its regulation of MAPK activity, not only plays a role in regulating PIGR in the context of proinflammatory signaling but is central to the regulation of constitutive (basal) PIGR expression.

### Independent validation the role of DUSP16 on the control of PIGR expression in epithelial cells

Although HT-29 cells are a widely used model for functional studies of intestinal epithelial cells, it is important to confirm that our observations are not limited to this cell line rather than being more broadly applicable to epithelial cells. We have therefore replicated our major findings related to the function of DUSP16 in the HCT-15 colonic epithelial cell line. Specifically, we increased the expression of DUSP16 in HCT-15 cells by lentiviral transduction of its ORF, in three independent infections as we had done in HT-29 cells. This transduction of the DUSP16 ORF not only led to the increased expression of DUSP16 in the HCT-15 cells, but also an important increase in the expression of PIGR, which was potentiated by treatment with TNF-alpha (**Additional file 2: Fig.S9**). In addition, the increased expression of DUSP16 led to a decrease in the steady-state phosphorylation levels of ERK, JNK, and p38 (**Additional file 2: Fig.S10A**). Finally, the expression of DUSP16 had a dramatic effect on JNK, and p38 phosphorylation levels in cells responding to TNF-alpha treatment (**Additional file 2: Fig.S10B**). These results are entirely concordant with those obtained in HT-29 cells.

## DISCUSSION

Genetic studies of IBD have met with significant success, with international collaborative studies identifying and validating over 200 genomic loci associated with CD, and/or UC, with most loci being shared by both. (1, 2, 13). For a limited number of these loci (e.g. IL23R, ATG16L1), the original GWAS studies identified the likely causal genetic variants (3, 4). Targeted sequencing identified novel variants in these same genes as well as identified IBD-associated coding variants in a handful of genes from loci containing multiple genes, thus prioritizing their corresponding genes as causal (5, 6, 54). High-density association mapping of GWAS loci pinpointed additional putative causal genes and their variants (7, 8). Despite these significant efforts, the causal gene(s) for the majority of GWAS loci remain to be identified. This highlights the need for developing novel approaches that are complementary to the genetic association studies, for example integration of eQTL datasets from relevant cell types, in order to identify causal genes within GWAS loci (62). Functional studies have provided confirmatory evidence as well as mechanistic insights for many of these genes (and variants) (4, 6, 9, 10, 17, 63). While the majority of functional studies have focused on immune cells, there is little doubt that the epithelial cell compartment has an important role to play in IBD, given its central position at the interface of the gut microflora and host immune system. Indeed, there is emerging evidence that genes involved in intestinal barrier function are implicated in common forms of IBD (12, 41, 64, 65) as well as in very early onset IBD (66, 67). Given this, we were interested in developing an approach to examine validated IBD loci for putative causal genes exerting their effect within epithelial cells. As opposed to RNAi-based screens that typically focus on the identification of genes that impact on a targeted set of known functions, we opted for an approach that involved increasing the expression of genes that already had a least a basal expression within epithelial tissues and cell lines and determining the impact of the gene expression modulation on the cell’s transcriptome, with the goal of identifying genes that impact on shared IBD-relevant epithelial functions. Just as for other functional genomic screens, this approach is not meant to mimic disease pathophysiology but rather to place genes in biological pathways in order to guide subsequent functional studies. We selected to perform this screen in the HT-29 cell line as it is a human intestinal epithelial cell line that is very frequently used for functional studies. We observed that the results from this approach were robust, and most informative for genes encoding transcription factors and proteins involved in signaling cascades, as opposed to structural proteins where there was much less of an impact on the transcriptome as could be expected.

Using a “similarity score” approach to cluster genes with shared effects on the transcriptome, we identified two large clusters of genes. Both of these clusters contained proven IBD causal genes, as well as genes from IBD loci where a causal gene has yet to be identified. Specifically, Cluster 1 contained the known IBD causal genes IFIH1, SBNO2, NFKB1 and NOD2, as well as genes from other IBD loci (ZFP36L1, IRF1, GIGYF1, OTUD3, AIRE and PITX1), whereas Cluster 2 contained the known causal gene KSR1 and implicated DUSP16 from another IBD locus. Analyses of the functional annotations of the transcripts most affected by the modulation of expression of these IBD genes suggested that Cluster 1 genes appear to be related to each other with respect to their involvement in the type I interferon response to viral components of the microbiome and/or the pro-inflammatory response to bacterial components of the microbiome. On the other hand, Cluster 2 genes appear to be implicated in epithelial structure and function, as well as mucosal defense, with the latter acting in part via the control of transcytosis of secretory immunoglobulins across enterocytes.

We therefore propose that multiple IBD genes play an important role within the intestinal epithelium in terms of regulation of the type I interferon response to viral pathogens and others in the transcriptional regulation of the anti-bacterial response. While for many of these genes (e.g. IFIH1, NFKB1, IRF1, NOD2, FOS) the association with anti-microbial functions has been well established, for others this association is novel. For example, OTUD3 is a known deubiquitylase of the tumour suppressor PTEN that has recently been shown to play a critical role in the antiviral response of innate immune cells and thus may play a similar role in intestinal epithelial cells (68, 69). ZFP36L1, on the other hand, is part of a family of RNA binding proteins (RBP) that bind A/U rich elements (ARE) in the 3’ untranslated region (3’UTR) of mRNAs and promotes of RNA decay. Most reports studying the role of this gene family have focused on immune cells and have proposed a role for ZFP36 in regulating pro-inflammatory responses (through the regulation of RNA levels of inflammatory cytokines) and a role for ZFP36L1/L2 in T- and B-lymphocyte development (70). Of relevance to the current study, a recent report using a non-immune model (Hela cells) has proposed that ZFP36 could regulate gene targets at the transcriptional level, rather than RNA stability, and has shown that ZFP36 expression leads to an increased expression of genes involved in Type I interferon signalling and antiviral response, similar to what we have observed for ZFP36L1 (71). GIGYF1, a GYF domain-containing scaffold protein, can promote translational repression of specific targets through its interaction with mRNA-binding proteins including ZFP36 family member TTP.

In terms of type I IFN response, it is clear that practically all cells within the body can mount an anti-viral response. Viruses are detected in a virus- and cell-specific manner through different sets of PAMPs, such as Toll-like receptors (TLRs), RIG-I-like receptors (RLRs) and NOD-like receptors (NLRs), inducing the expression of Type I IFNs. Type I IFNs can then activate, in an autocrine and paracrine fashion, the transcription of interferon response genes (IRGs), including several different anti-viral defense factors, and induce a refractory anti-viral state. Intestinal epithelial cells (IECs) play a crucial role in intestinal barrier functions, contributing multiple protective mechanisms to avoid the penetration of pathogens from the outside environment into the body including Type I IFN and Type III responses. Type I IFNs mainly include IFN-α and IFN-β and interact with the heterodimeric receptor IFNAR (72, 73). Type III IFNs include four IFNλ molecules whose receptor is composed of the IL-28Rα/IL-10R2 complex, which is specifically expressed on epithelial cells (74). While the role of Type I IFN response in protecting IECs against different types of enteric viruses (rotavirus, reovirus and norovirus) has been well described, their role in controlling intestinal homeostasis and inflammation is not as well characterized, with contradictory results in the context of DSS-induced colitis (75, 76). While the results presented herein make a strong case for multiple IBD genes being involved in the Type I IFN response, it is possible that they may also act on Type III signaling as the HITS for these genes were also enriched for gene annotations “Type III interferon signaling” and/or “response to interferon-gamma”, although to a lesser extent (**Additional file 3: Cluster 1 data**). While dysregulated Type I interferon signaling has typically been associated with autoimmune diseases such as systemic lupus erythematosus in the context of circulating immune cells (77, 78), given the unique position of IEC at the interface with enteric viruses, it is conceivable that dysregulated interferon signaling within IEC may not only impact the epithelium but the immune cells that are in close proximity. In this context, it is interesting to note that while most of the genes in Custer 1 are fairly ubiquitously expressed, OTUD3 and PITX1 are primarily expressed in intestinal cells amongst the tissues and cell lines tested.

Cluster 2 was centered around the known IBD gene KSR1 that acts as a scaffold for the Ras/Raf/MAPK signaling pathway (56). The data presented herein suggests that KSR1 is involved in regulating a broad set of intestinal functions and cell differentiation pathways. DUSP16 was the closest gene to KSR1 in the similarity score and shared multiple functions and top HITS, highlighting potential roles in dietary metabolism, homeostasis and defense. Our follow-up studies were aimed at obtaining better understanding the function of DUSP16 in intestinal epithelial cells, with a particular focus on the transcriptional control of PIGR as the latter’s expression is enriched in gastrointestinal mucosa and has a known role in the protection of the mucosa from intestinal microbiota (79). Specifically, pIgR is expressed at the basolateral surface of epithelial cells where it binds polymeric immunoglobulin molecules (IgA and IgM). The complex is then internalized and transported across the cell, via transcytosis, to the apical surface where Ig molecules are secreted as sIg (IgA/M bound to SC) following the cleavage of part of the pIgR. sIg can then modulate microbial communities and pathogenic microbes via several mechanisms: agglutination and exclusion from the epithelial surface, neutralization, or via host immunity and complement activation (79, 80). In addition, it has also been suggested that pIgR-bound Ig molecules can also play a role in shuttling foreign antigens from the basolateral to the luminal compartment via transcytosis, thus protecting the integrity of the epithelial barrier (79, 81, 82).

It has previously been shown in fibroblastic cell lines (COS-7 and NIH3T3) that DUSP16 has dual specificity protein phosphatase activity and functions to regulate MAP kinases (57, 58). We were able to demonstrate that this was also the case in human epithelial cells, as well as demonstrate its specificity for JNK and p38, but to a much lesser extent ERK, which is consistent with published studies (57, 58). Moreover, we were able to demonstrate that inhibition of p38 and ERK also increased the basal expression of PIGR. As it had been previously reported that TNF-α can regulate the expression of PIGR (60, 61), we examined the impact of DUSP16 and found that it potentiated the effect of TNF. While a functional link between DUSP16 and PIGR regulation can clearly be defined via its role in inhibiting MAPK activity, such a link to KSR1 may not be as direct. As stated above, KSR1 acts as a scaffold for the cytoplasmic Ras/Raf/MAPK (ERK) signaling pathway in response to EGF and has been suggested to carry a kinase activity. As a scaffold protein, KSR1 has the capacity to redirect ERK from the cytosol to cytoplasmic membrane; expression of KSR1 may therefore act to sequester a larger proportion of the cell’s ERK pool to the cytoplasmic membrane, in effect leading to an overall reduction of nuclear ERK activity. In that regard, expression of KSR1 would lead to a similar effect to DUSP16 on ERK activity but through an unrelated mechanism. While this mechanism of action of KSR1 in our model is purely speculative, it may warrant further investigation.

We therefore propose a model by which lower levels/activity of KSR1 or DUSP16 would lead to increased nuclear MAPK activity (ERK and p38) which could contribute to disease susceptibility, in part, by interfering with PIGR expression and thus limiting the transport of IgA antibodies across IECs and diminishing the ability of the mucosal barrier to protect against intestinal microbiota. Interestingly, PIGR resides within one of the 163 IBD-associated regions identified by GWAS, although outside the boundaries defined by the current project; in fact, this region contains many interesting functional candidate genes for IBD, including the candidate causal gene IL10. Moreover, two recent studies of inflamed intestinal tissue from patients with UC observed that the extensive remodeling of inflamed tissue is associated with somatic mutations that converge on a handful of genes, including PIGR, implicated in the downregulation of IL-17 signalling and other pro-inflammatory signals (83, 84).

As with all large-scale genomic screens, it was important to validate the results of our transcriptomic screen in HT-29 cells in order to ensure that our observations were not simply a result of the experimental design or limited to the cell line used. In the current study this was achieved by (1) analysis of independent datasets to provide bioinformatic support for the HNF4-alpha, SMAD3/SMAD7, and IFIH1 genes; (2) studying the impact of disease-associated coding variants in IFIH1; (3) using complementary genetic modulation approaches for DUSP16 (stable, inducible, and knockdown); (4) using pharmacologic perturbations to validate DUSP16’s biological pathway; and (5) validation of key findings in the independent intestinal epithelial cell line HCT-15. Nonetheless, future studies will be required to deepen our functional understanding of how these various genes play a role in IBD pathophysiology, as well as follow-up on the observations reported herein for the genes that we have yet to perform validation experiments.

## CONCLUSIONS

In conclusion, employing an analysis strategy that is analogous to the “guilt by association” approach whereby genes that have correlated patterns of expression within an expression network display increased likelihood of having shared functions (85), the current transcriptomic-based screen in the HT-29 human colonic epithelial cell line has highlighted how specific IBD genes are potentially functionally related to each other, based on their shared impacts on the cell’s transcriptome following a lentiviral-mediated expression of the relevant ORFs. This has enabled the prioritization of likely causal genes within multi-gene loci without a previously identified causal gene. Taken together, the results highlight that multiple IBD genes are involved in a variety of host defense mechanisms operating within epithelial cells. Moreover, these pathophysiologic mechanisms extend beyond the classical physical barrier that epithelium provides and beyond the cells specialized in host defense, such as Paneth cells, that have previously been implicated by previous genetic studies of IBD (1, 12, 17, 41, 64, 65, 86). Moreover, this adds to the growing evidence in support of epithelial cells playing key roles in the local inflammatory processes (87–89).

## DECLARATIONS

### Ethics approval and consent to participate

Not applicable

### Consent for publication

Not applicable

### Availability of data and materials

The datasets used and/or analysed during the current study are available from the corresponding author on reasonable request.

### Authors’ contributions

Overall project and/or experiments conceived by JDR, RJX, GC and PG with input from the iGenoMed Consortium members. Laboratory work and analyses were supervised by GC, PG and JDR. Agilent chip design was performed by SF, GC and PG. Data generation performed by AA, CB, GL, HG, JCN, JP, MJD, NM, SD. Bioinformatics analyses, and management of datasets and samples, were performed by GB, FL and SF. Statistical analyses were performed by GB. Interpretation of results by GB, JCN, PG, and JDR. The manuscript was written by PG, GB, and JDR with contributions from JP and JCN. Overall study supervision and management performed by JDR. All authors read and approved the final manuscript.

## Acknowledgements.

We would like to thank Drs. David Root (Broad Institute) and Gerald T. Nepom (Benaroya Research Institute) for their helpful discussions. We would like to thank the Array Technologies Unit at the Génome Quebec McGill Innovation Centre for processing the Illumina Human HT-12 v4 beadchips. We would like thank members of the Laboratory for Genetics and Genomic Medicine, in particular Julie Thompson Legault for her help in preparation of the manuscript as well as Mélanie Burnette, Jessica Desjardins and Marie-Pier Mathieu for technical assistance.

The members of the iGenoMed Consortium at the time of this study were (in alphabetical order): Alain Bitton^1*^, Gabrielle Boucher^2^, Guy Charron^2^, Christine Des Rosiers^2,3*^, Anik Forest^2^, Philippe Goyette^2^, Sabine Ivison^4^, Lawrence Joseph^5*^, Rita Kohen^1^, Jean Lachaine^6*^, Sylvie Lesage^3,7*^, Megan K. Levings^4*^, John D. Rioux^2,3*^, Julie Thompson Legault^2^, Luc Vachon^8^, Sophie Veilleux^9*^, Brian White-Guay^3*^. Affiliations: ^1^McGill University Health Centre, Montreal, Quebec; ^2^Montreal Heart Institute Research Center, Montreal, Quebec; ^3^Université de Montréal, Faculté de Médecine, Montreal; ^4^University of British Columbia, Vancouver; ^5^McGill University, Faculty of Medicine, Department of Epidemiology, Biostatistics and Occupational Health; ^6^Université de Montréal, Faculté de Pharmacie; ^7^Maisonneuve-Rosemont Hospital, Research Center, Montreal; ^8^LV Consulting, Montreal; ^9^Université de Laval, Québec. *Principal Investigators on grant # GPH-129341 with JDR as Leader and AB as co-Leader.

## SUPPLEMENTARY INFORMATION

Additional file 1: Additional_File_1.xlsx; Supplementary Tables S1-S12

Additional file 2: Additional_File_2.pdf; Supplementary Figures S1-S12, Box1 & Box 2

Additional file 3: Additional_File_3.xlsx; Supporting data for Cluster 1 genes (Complete list of HITs and functional enrichment results)

Additional file 4: Additional_File_4.xlsx; Supporting data for Cluster 2 genes (Complete list of HITs and functional enrichment results)

