## Additional File 2 for "Functional screen of Inflammatory bowel disease genes reveals key epithelial functions"

**Index**

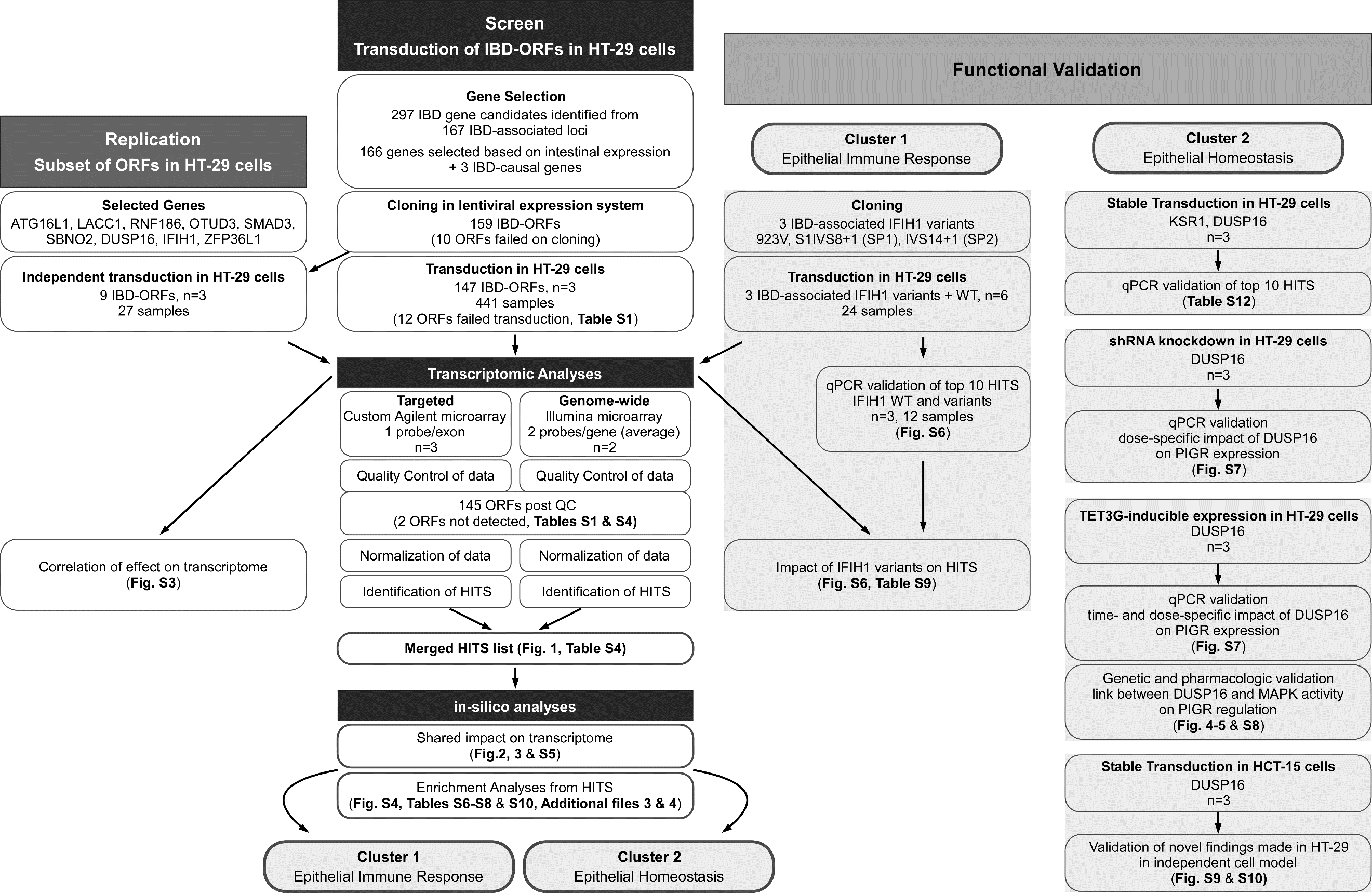

**Fig.S1: Flow chart of the expression-based functional screen, with replication and validation steps.** A step-by-step overview illustrating the different phases of our expression-based functional screen is presented, with emphasis on selection of IBD gene candidates, transcriptomic analysis, replication and validation studies. Further details for each stage can be found in the Materials and Methods section, as well as in the different display items referenced herein.

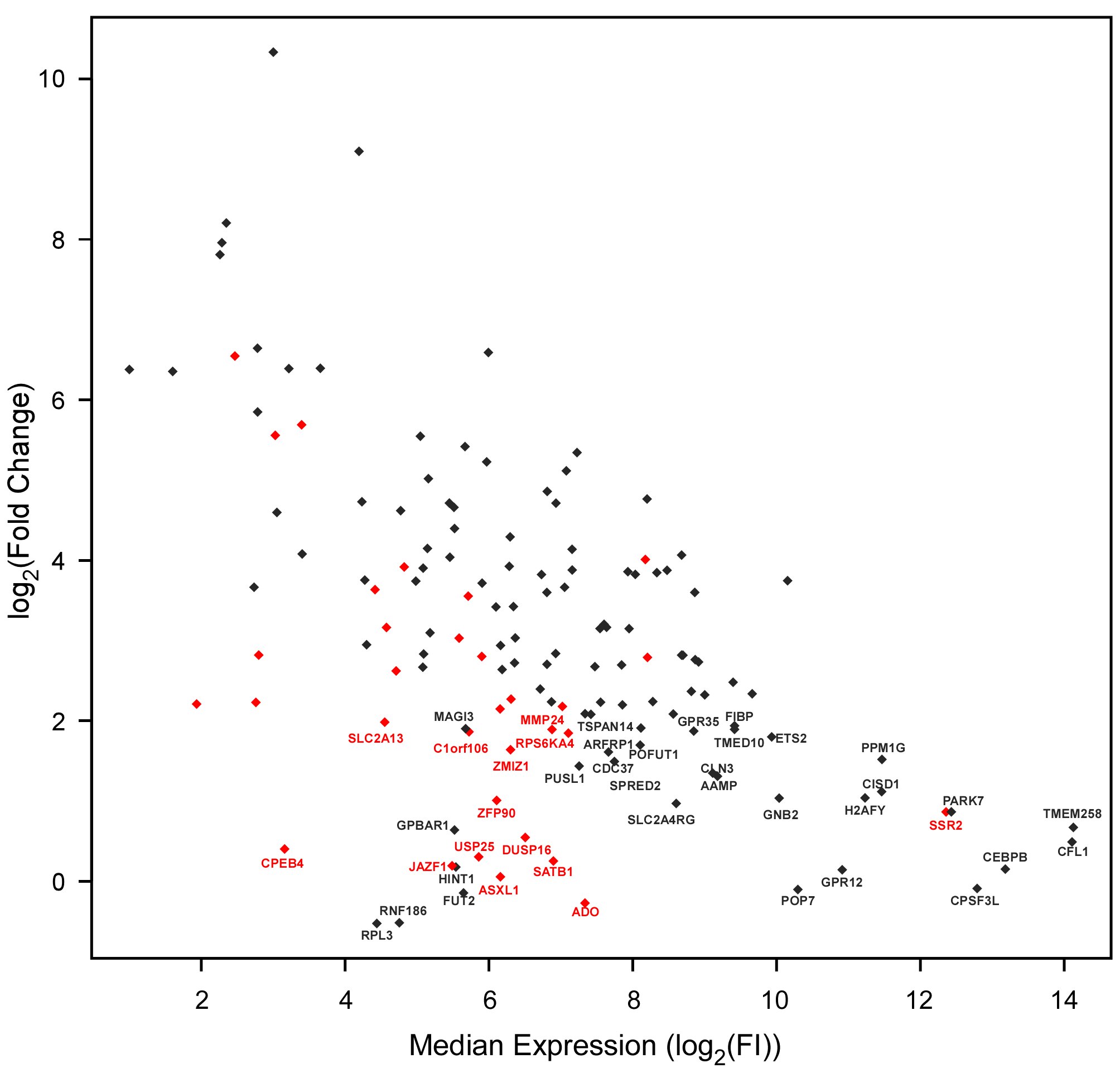
**Fig.S2: Fold induction of each IBD-gene ORF following their transduction in HT-29 cells.** Expression level for each of the different IBD genes was evaluated in the transcriptomic data from the Illumina Human HT-12 v4 Beadchip whole genome microarray and plotted as a fold-change (y-axis) against its median endogenous expression (x-axis) in HT-29 cells. The cDNA sequence of the ORFs in red were synthesized as optimized sequences and as such are often not well detected by probes found on expression microarrays designed to detect endogenous gene sequences. Confirmation of increased expression for ORFs not showing a clear increase from microarray data was performed via endpoint PCR (see **Additional file 1: Table S4**).

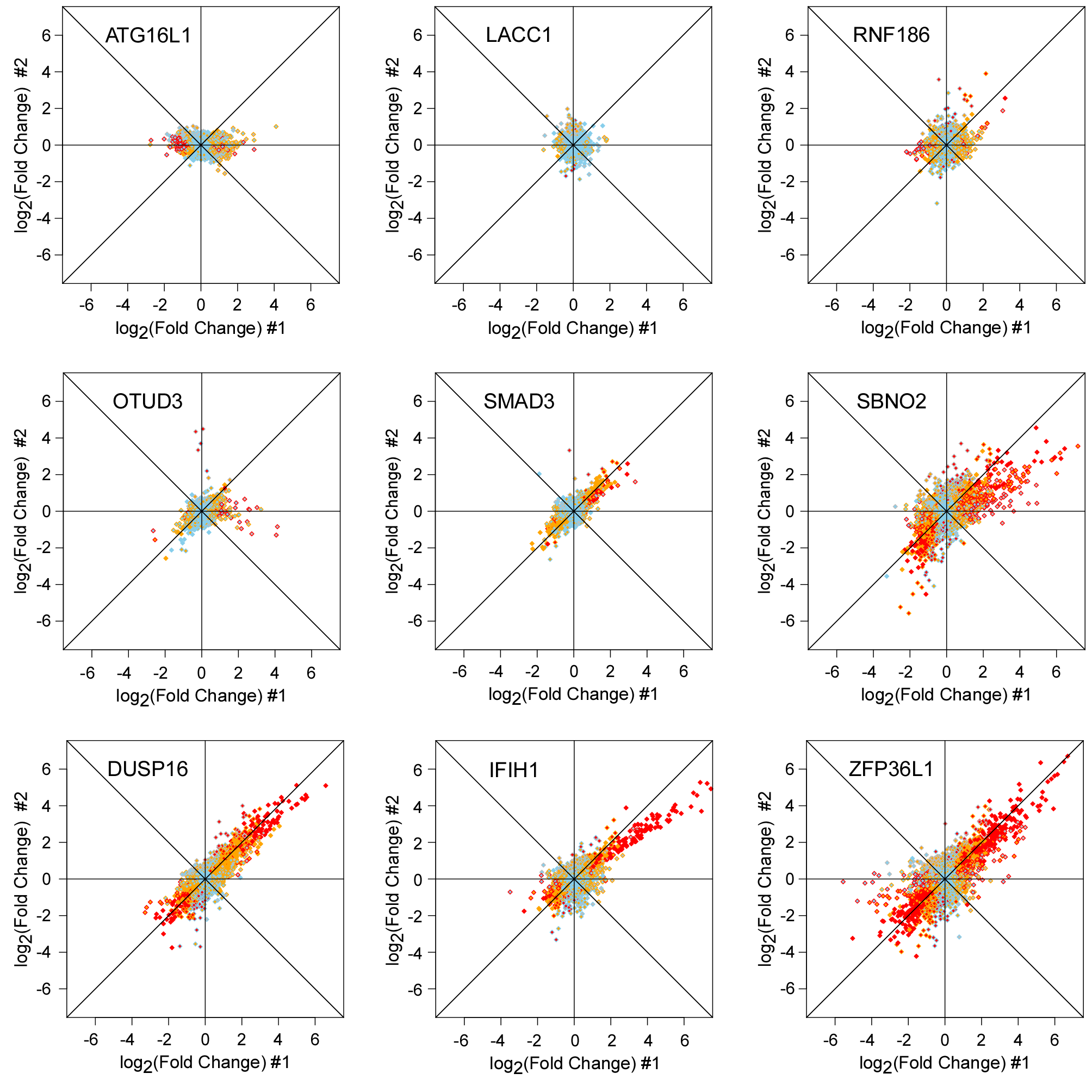

**Fig.S3: Correlation of effect of independent sets of replicated ORFs on HT-29 transcriptome.** As a validation of the effects observed on the transcriptome following ORF transduction, a set of nine ORFs were repeated, each with an additional set of three independent transduction experiments performed 4, 5, 17, 10, 18, 15, 18, and 11 months apart for ATG16L1, LACC1, RNF186, OTUD3, SMAD3, SBNO2, DUSP16, IFIH1 and ZFP36L1, respectively. Variation between sets of replicates includes effect of independent infection dates, RNA extraction, expression arrays and batches. Each dot represents a single detectable probe from the genome-wide array tagging a specific gene in the HT-29 transcriptome (see **Fig.1**). The x-axis and y-axis show the effect of two different ORFs on the transcriptome, as the log2-transformed fold-induction compared to baseline. Skyblue are probes with expression value within expected variation (|Z| ≤ 2), orange represent probes suggestively outside the range (|Z|>2 ) and red represent probes outside expected range of variation (|Z| > 4), and gray are probes with expression value below our detection threshold; this color code is applied to the inside of dots for x-axis data and the border of dots for y-axis data.

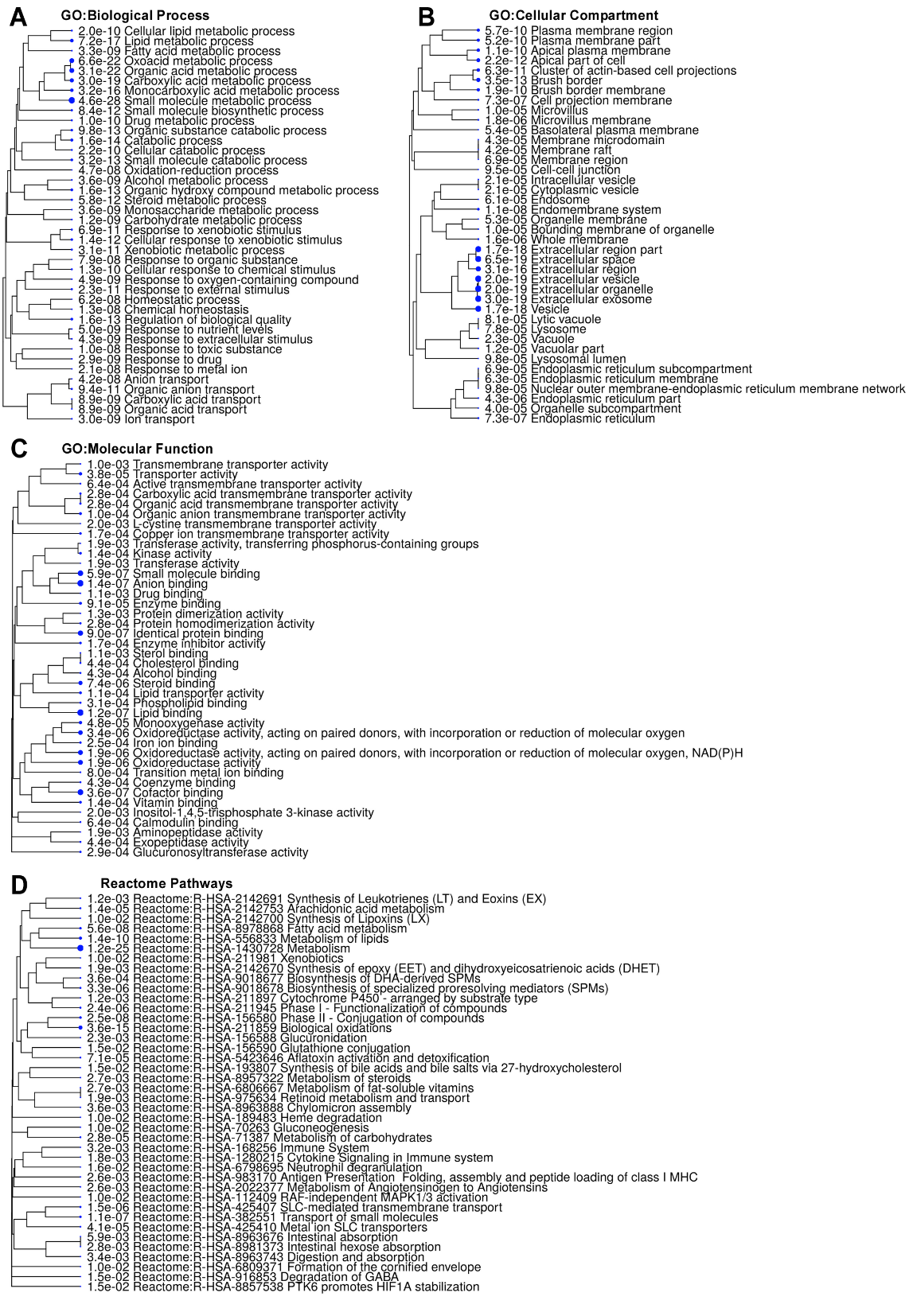

**Fig.S4: Functional enrichment analysis of HNF4A HITS.** Enrichment for Gene Ontology (GO) terms and biological pathways was performed in ShinyGO v0.61, an online gene ontology enrichment analysis tool (PMID: 31882993), using up-regulated HITs obtained from the transduction of the ORF for HNF4A. Hierarchical clustering trees constructed based on the correlation (many shared genes) among the top 40 most significant functional enrichments for each category are shown (at a P-value threshold of <0.05). Dot size is proportional to significance of P-values.

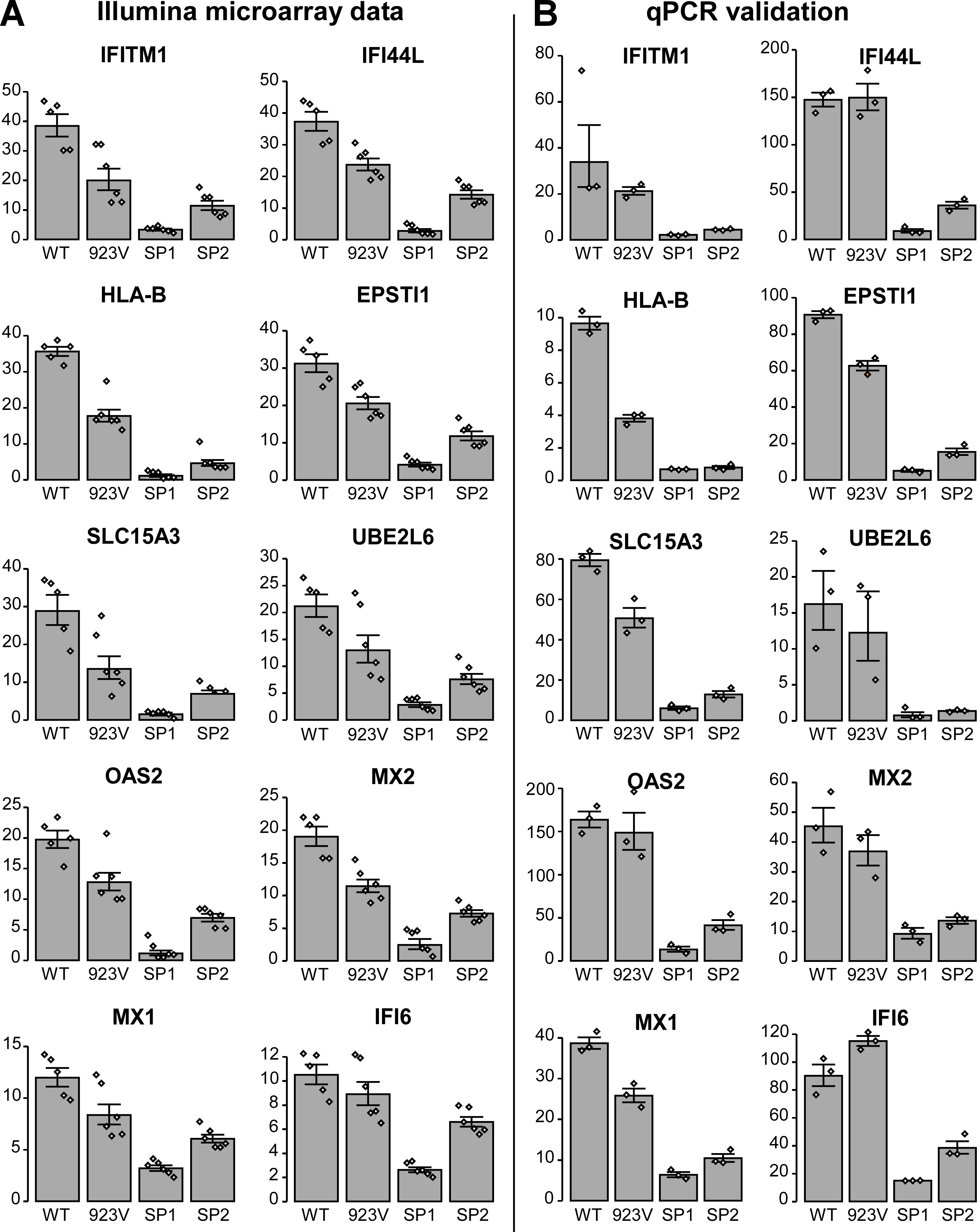

**Fig. S5: Impact of IBD-associated IFIH1 variant on the top 10 IFIH1 HITS. (A)** The effect of wild-type IFIH1 (n=5) and three of its IBD-associated non-synonymous coding variants (923V, IVS8+1 (SP1) and IVS14+1 (SP2); n=6 for each) on the induction of the top 10 IFIH1 HITS in our HT-29 transcriptomic dataset is shown. The values shown were extracted from the same dataset used in **Fig.3B**; for the effect of IFIH1 variants on all IFIH1 HITs, see **Additional file 1: Table S9**. Fold changes in expression compared to the baseline, with standard error are plotted (y-axis); the values for independent replicates are shown (grey lozenges). **(B)** The results presented in panel A were validated by qPCR analysis in a subset of samples (n=3). Fold changes in expression compared to the empty vector, with standard error are plotted (y-axis); the values for independent replicates are shown (grey lozenges).

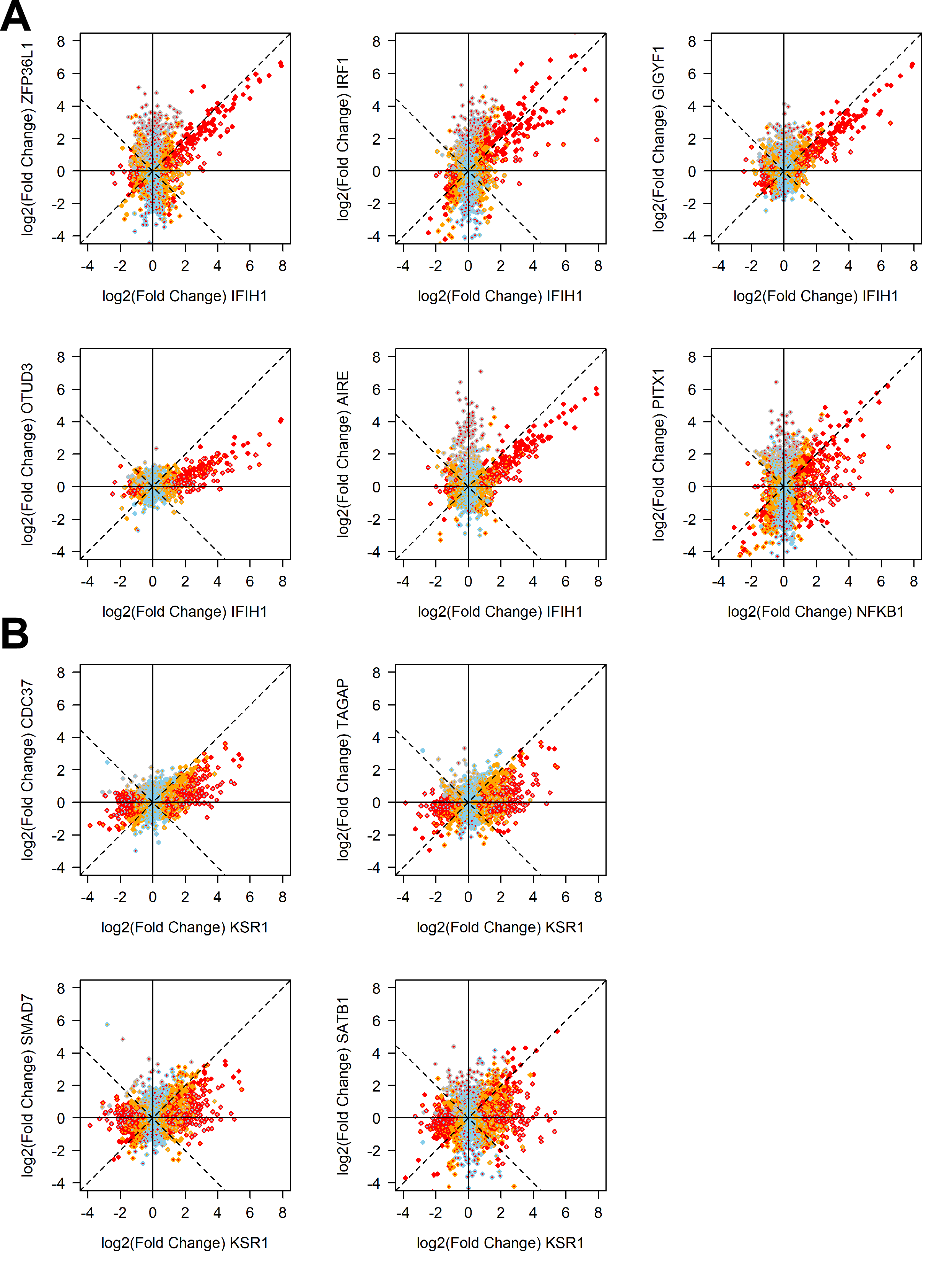

**Fig.S6: Shared effects of IBD causal genes IFIH1, NFKB1 and KSR1 to IBD gene candidates ORF neighbors in cluster 1 and 2.** Correlation plot of the effect of different IBD causal genes against different IBD gene candidate cluster neighbors are shown for **(A)** cluster 1 ORFs and **(B)** cluster 2 ORFs. Selection of comparison pairs was made based on proximity in similarity plot (Fig.2), as well as on number of shared HITs and shared functional enrichments. Each dot represents a single detectable probe from the genome-wide array tagging a specific gene in the HT-29 transcriptome (see **Fig.1**). The x-axis and y-axis show the effect of two different ORFs on the transcriptome, as the log2-transformed fold-induction compared to baseline. Skyblue are probes with expression value within expected variation (|Z| ≤ 2), orange represent probes suggestively outside the range (|Z|>2 ) and red represent probes outside expected range of variation (|Z| > 4), and gray are probes with expression value below our detection threshold; this color code is applied to the inside of dots for x-axis data and the border of dots for y-axis data.

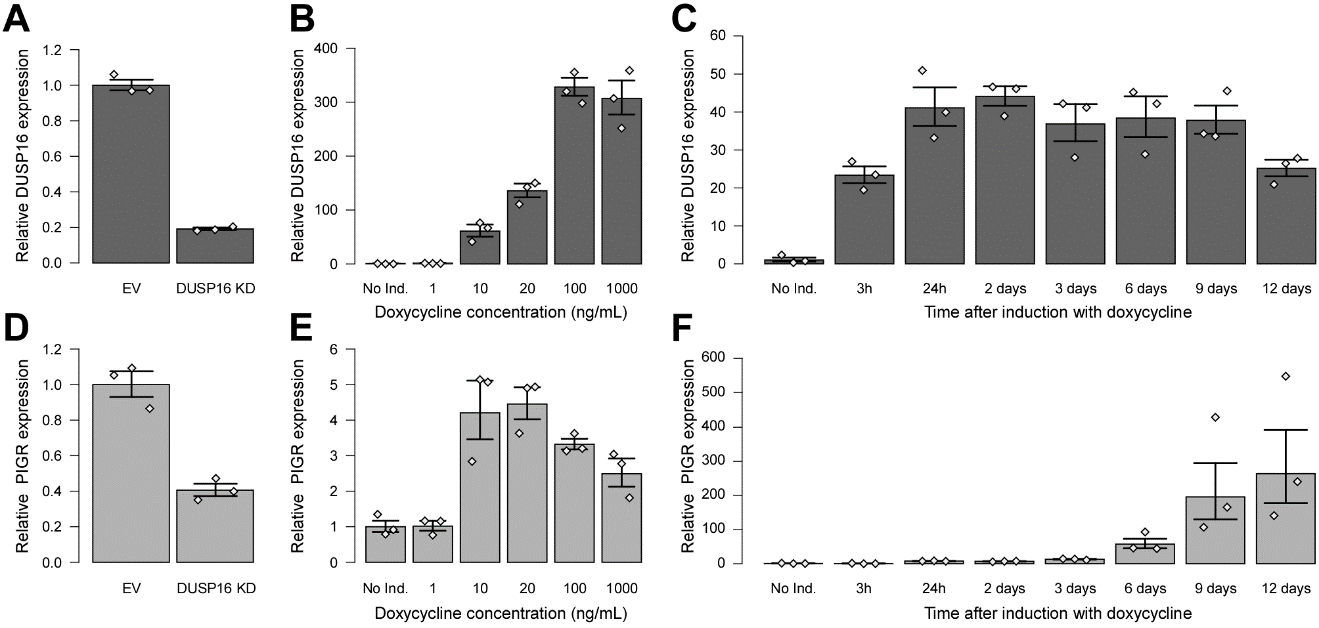

**Fig.S7: Impact of modulating DUSP16 levels on PIGR expression.** The impact of modulating DUSP16 expression levels in HT-29 cells, either via shRNA knock-down or ORF induction, on PIGR mRNA levels was evaluated. (**A, D**) HT-29 cell lines stably transduced with lentiviral shRNA expression vectors, either empty vector (EV, n=3) or containing a DUSP16-specific shRNA (DUSP16 KD, n=3) (TRCN0000052017) were evaluated for endogenous DUSP16 **(A)** and PIGR **(D)** mRNA expression levels; qPCR results from the replicates were combined and mean expression values relative to the cells expressing the empty vector are shown. (**B, C, E, F**) Exponentially growing HT29-pLVXEE1α-Tet3G cell lines stably transduced with the TET3G-inducible expression plasmid for DUSP16 ORF (n=3) were left untreated (N.T.) or stimulated with either increasing doses of Doxycycline for 24 hours (dose response (**B, E**)) or with Doxycycline at a concentration of 10 ng/ml for different times (time course (**C, F**)) to induce DUSP16 expression. Expression levels of DUSP16 (**B, C**) and PIGR (**E, F**) mRNA were evaluated; qPCR results from the replicates were combined and mean expression values relative to samples without treatment with doxycycline (N.T.) are shown. All results are shown as geometric means with SEM. The values for independent replicates are shown (grey lozenges).

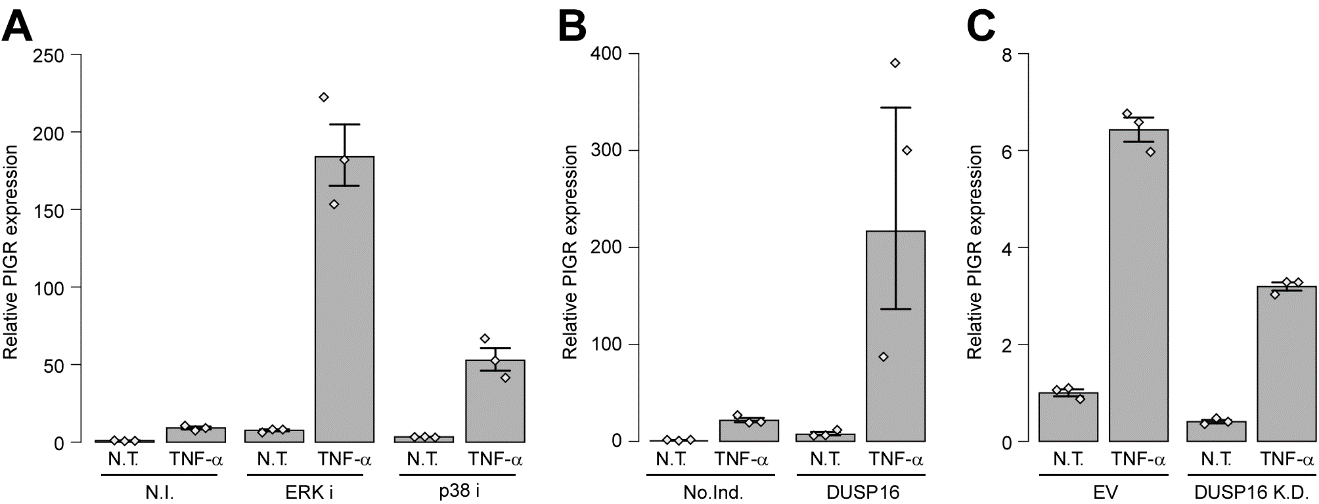

**Fig.S8: Impact of MAPKs inhibition on PIGR expression in response to TNF-α in HT-29 cells.** The impact of an inhibition of ERK or p38 activity, either chemically or following induction of DUSP16 expression, on PIGR mRNA levels in response to TNF-α in HT-29 cells was evaluated by qPCR. **(A)** Exponentially growing HT29-pLVX-Tet3G cell lines (n=3) were either left untreated (no inhibitor; N.I.) or treated with chemical inhibitors specific for ERK (PD98059, ERKi) or p38 (SB203580) at a concentration of 10uM for 24 hrs and then either left untreated (N.T.) or stimulated with 10 ng/ml TNF-α for 3 hrs before total RNA was isolated. Expression levels of PIGR were evaluated via qPCR; the qPCR results from the replicates were combined and mean expression values relative to samples without inhibitors (N.I.) or treatment (N.T.) are shown. **(B)** Exponentially growing HT29-pLVX-Tet3G cell lines stably transduced with the TET3G-inducible expression plasmid for DUSP16 ORF (n=3) were either left untreated (No.Ind.) or stimulated with Doxycycline at a concentration of 10 ng/ml for 24 hrs to induce DUSP16 expression and then either left untreated (N.T.) or stimulated with 10 ng/ml TNF-α for 3 hrs. Expression levels of PIGR were evaluated via qPCR; the qPCR results from the replicates were combined and mean expression values relative to samples without inhibitors (N.I.) or treatment (N.T.) are shown. **(C)** HT-29 cell lines stably transduced with lentiviral shRNA expression vectors, either empty (EV, n=3) or containing a DUSP16-specific shRNA (DUSP16 KD, n=3) (TRCN0000052017) were either left untreated (N.T.) or stimulated with 10 ng/ml TNF-α for 3 hrs. Expression levels of PIGR were evaluated via qPCR; the qPCR results from the replicates were combined and mean expression values relative to EV samples without treatment (N.T.) are shown. All results are shown as geometric means with SEM. The values for independent replicates are shown (grey lozenges).

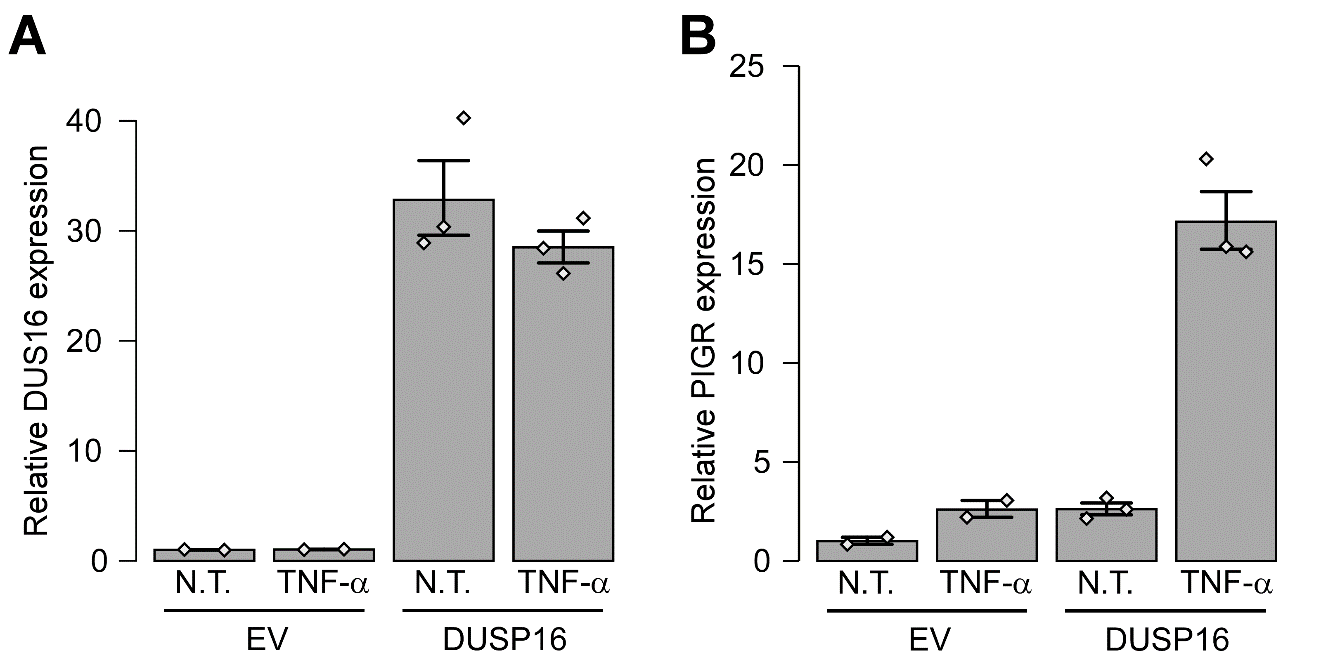

**Fig. S9: Impact of DUSP16 on PIGR expression levels in response to TNF-α in HCT-15 cells.** The impact of an increase in DUSP16 expression on PIGR mRNA levels in response to TNF-α was evaluated by qPCR in HCT-15 cells. Exponentially growing HCT-15 cell lines stably transduced with either empty pLVX-EF1a-IRES-PURO/eGFP lentiviral vector (EV, n=2) or containing the ORF encoding for DUSP16 (n=3) were either left untreated (N.T.) or stimulated with 10 ng/ml TNF-α for 3 hrs before total RNA was isolated. Expression levels of **(A)** DUSP16 and **(B)** PIGR were evaluated via qPCR; the qPCR results from the replicates were combined and mean expression values relative to samples without treatment (N.T.) are plotted. All results are shown as geometric means with SEM and the values for independent replicates are shown (grey lozenges).

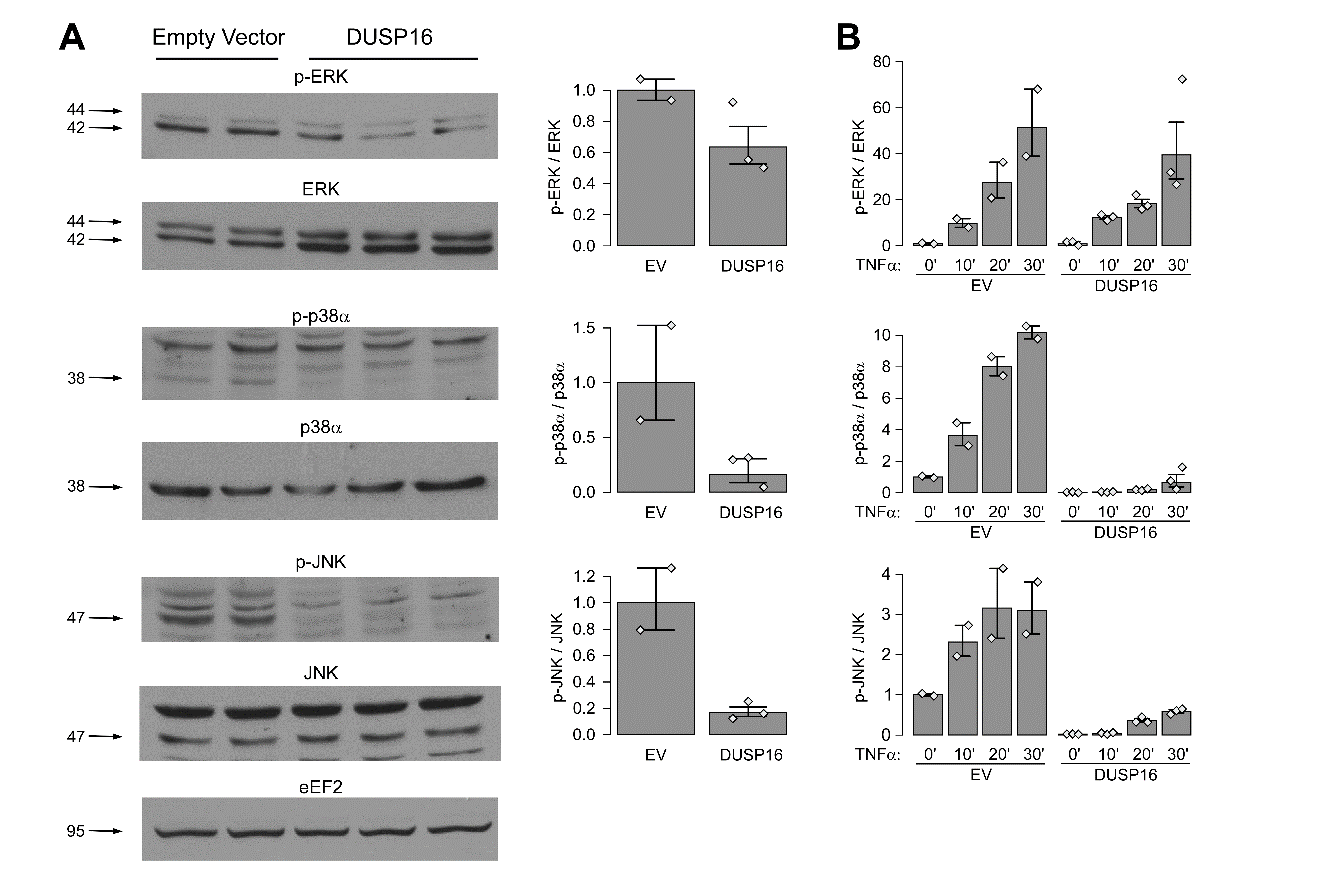

**Fig. S10: Impact of DUSP16 ORF on MAPKs phosphorylation levels in HCT-15 cells.** The impact of DUSP16 of an increase in DUSP16 expression on ERK, p38 and JNK phosphorylation levels (p-ERK, p-JNK and p-p38 respectively) was evaluated by Western blot analysis, in HCT-15 cells stably expressing DUSP16, using 2 different approaches. **(A)** The impacts of DUSP16 on steady-state MAPKs phosphorylation levels was measured. Exponentially growing HCT-15 cells stably transduced with either empty pLVX-EF1a-IRES-PURO/eGFP lentiviral vector (EV, n=2) or containing a codon optimized DUSP16 ORF (n=3) were harvested and whole-cell lysates obtained for the evaluation of the native and phosphorylated states of MAPKs by Western blot analysis. Cropped western blots are shown (left) and the level of phosphorylation of each MAPK, expressed as the ratio of phosphorylated MAPK over total MAPK (both corrected for the EEF2 loading control), is summarized in a graphical format (right) combining the replicates. A single representative gel is shown for EEF2 loading control; Full blots along with their respective EEF2 results for each primary target antibody tested are included in **Fig. S12, panel A**. **(B)** The impact of DUSP16 on the induction of MAPKs phosphorylation following serum starvation is measured. Using the same models as above, exponentially growing cells were first serum-starved for 24 hours (McCoy's 5A (Modified) medium, 0% FBS). The cells were then stimulated with 10 ng/ml TNF-α for different times to induce MAPK phosphorylation and whole cell lysates harvested for the evaluation of the native and phosphorylated states of MAPKs by Western blot analysis. A graphical representation of the phosphorylation state of each MAPK, expressed as the ratio of phosphorylated MAPK over total MAPK (both corrected for the EEF2 loading control), is shown combining the replicates. All results are shown as geometric means with SEM. The y-axis shows expression relative to the control condition (EV in **panel** **A** and EV 0’ in **panel** **B**). The values for independent replicates are shown (grey lozenges). Full blots for each replicate and all timepoints, along with EEF2 results for each primary target antibody tested are included in **Fig. S12, panel B-C**.

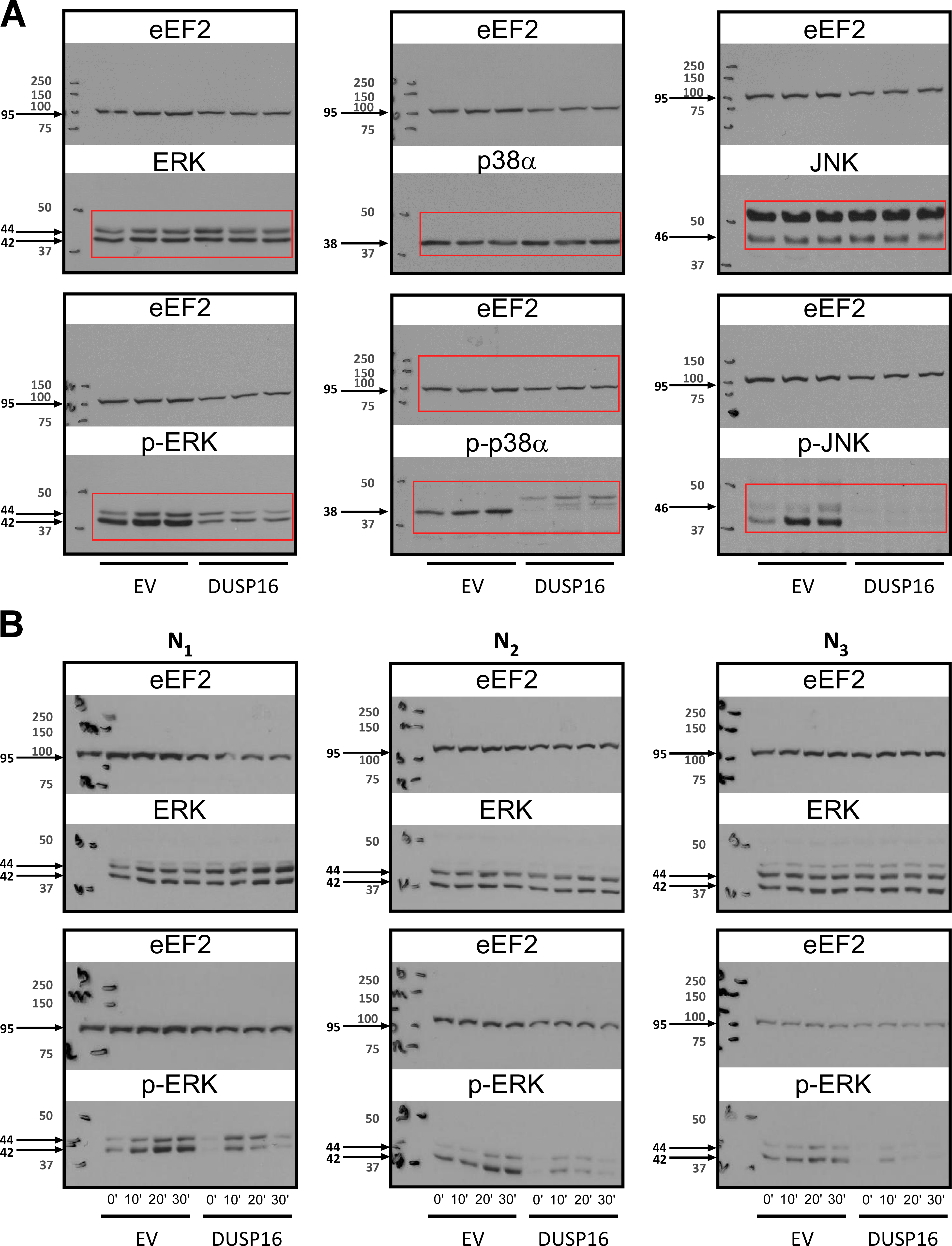

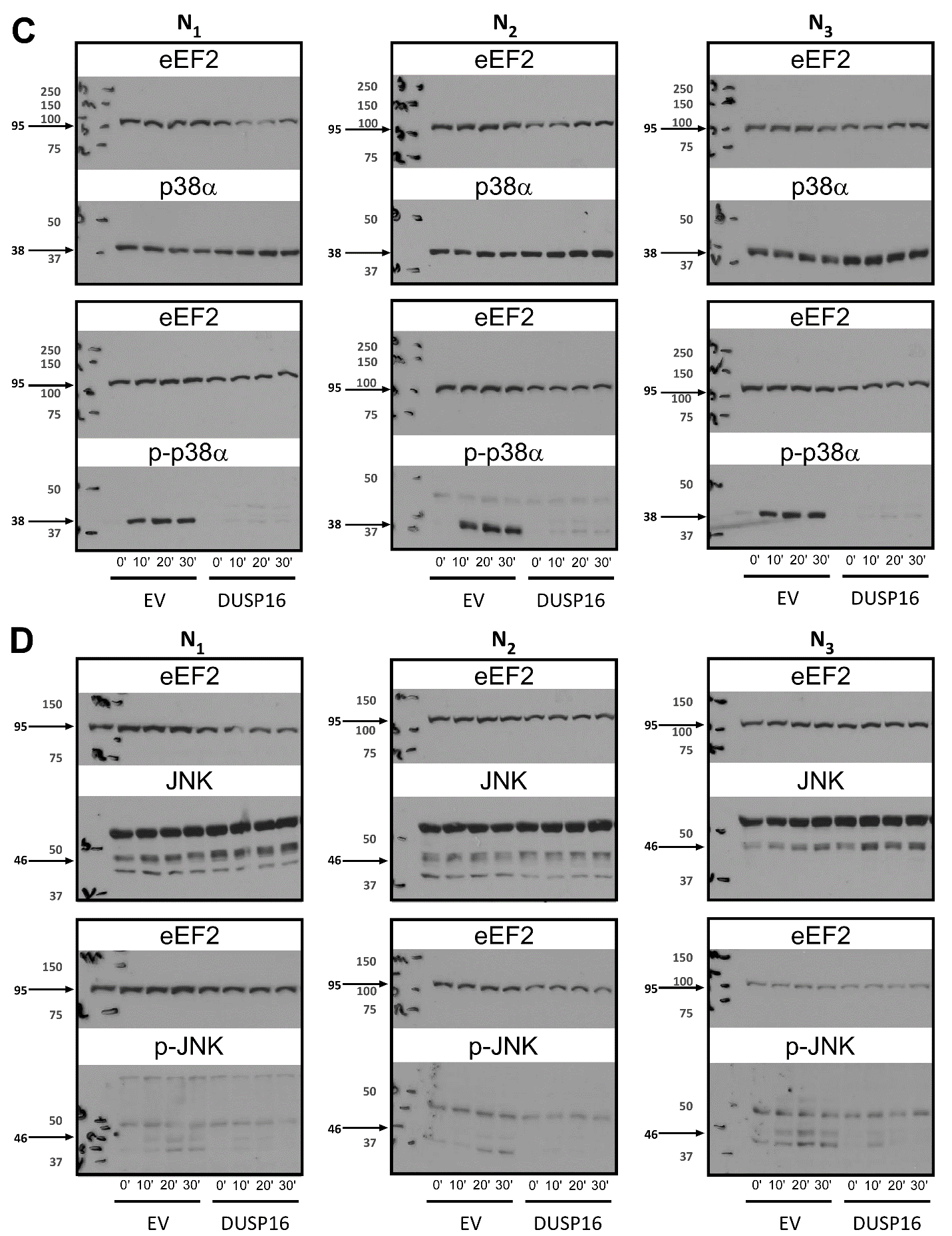

**Fig. S11: Western blots analysis (full immunoblots) of the impact of DUSP16 expression on MAPKs phosphorylation levels in HT-29 cells (all replicates used to generate Fig. 4A&B).**

**(A)** **Impact of DUSP16 on steady-state MAPKs phosphorylation levels**. Exponentially growing HT29-pLVX-Tet3G cells stably transduced with a TET3G-inducible expression plasmids, either empty (empty vector, EV) or containing the codon-optimized DUSP16 ORF (DUSP16), were stimulated with Doxycycline (1ug/ml) for 24 hours to induce DUSP16 expression and whole cell lysates were then harvested for the evaluation of the native and phosphorylated states of ERK, p38 and JNK by Western blot analysis. Whole cell lysates were run on SDS-PAGE and transferred onto nitrocellulose membranes (BioRad). The membranes were then cut in half at the level 75kDa to allow independent immunoblotting and detection of proteins of different sizes (primary test targets (MAPKs) and loading control (eEF2)); full immunoblots representing the two halves of the same membrane are shown boxed together. Region of immunoblots boxed in red are those shown in **Fig. 4A**. Molecular weight markers were run on each gel and are marked on images with corresponding sizes on the left; expected molecular weights of target proteins are indicated with arrows. Following detection, immunoblots were scanned into JPEG files and expression levels were determined using the Image J software (Fiji). Levels of each MAPK or p-MAPK were first corrected with their corresponding eEF2 loading controls, and then the level of phosphorylation of each MAPK, expressed as the ratio of phosphorylated MAPK over total MAPK, was calculated. The results are summarized in **Fig. 4A** combining the replicates (n=3).

**(B-C-D)** **Impact of DUSP16 on the induction of MAPKs phosphorylation following serum starvation**. Using the same cell models as above, exponentially growing cells were first serum-starved for 24 hours (McCoy's 5A (Modified) medium, 0% FBS) in the presence of the Doxycycline (1ug/ml) inducer. The cells were then stimulated with 10 ng/ml TNF-α for different times to induce MAPK phosphorylation and whole cell lysates harvested for the evaluation of the native and phosphorylated states of MAPKs by Western blot analysis (as described above); immunoblots for each MAPK and p-MAPK across three replicates (N_1_, N_2_, N_3_) are shown separately for **(B)** ERK, **(C)** p38, and **(D)** JNK detection. Levels of each MAPK or p-MAPK were first corrected with their corresponding eEF2 loading controls, and then the level of phosphorylation of each MAPK, expressed as the ratio of phosphorylated MAPK over total MAPK, was calculated. The induction of phosphorylation at different time points is shown in **Fig. 4B** (combining the three replicates) for the cells containing empty vector (EV (+)) and the DUSP16 ORF stimulated with Doxycycline (DUSP16 (+)).

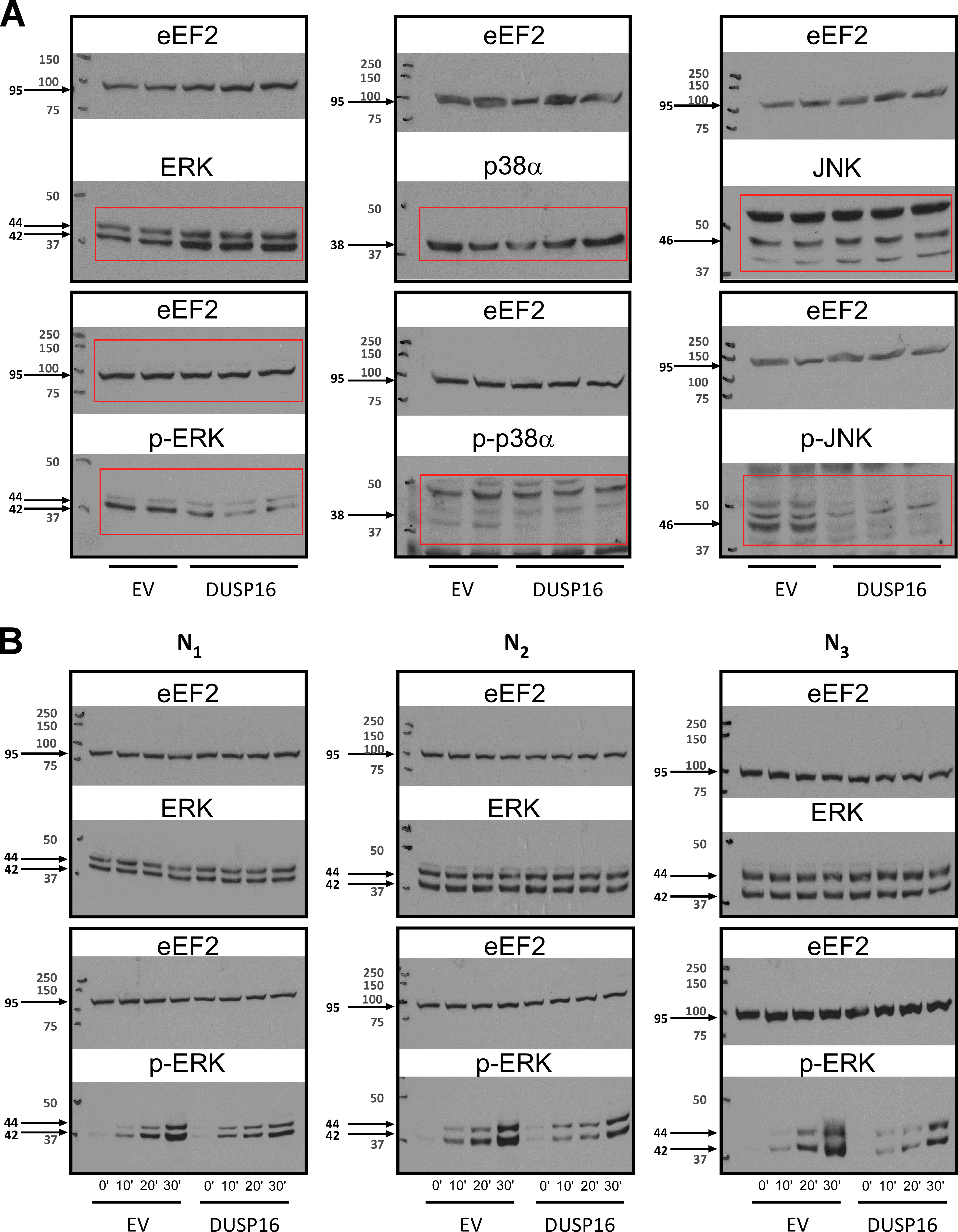

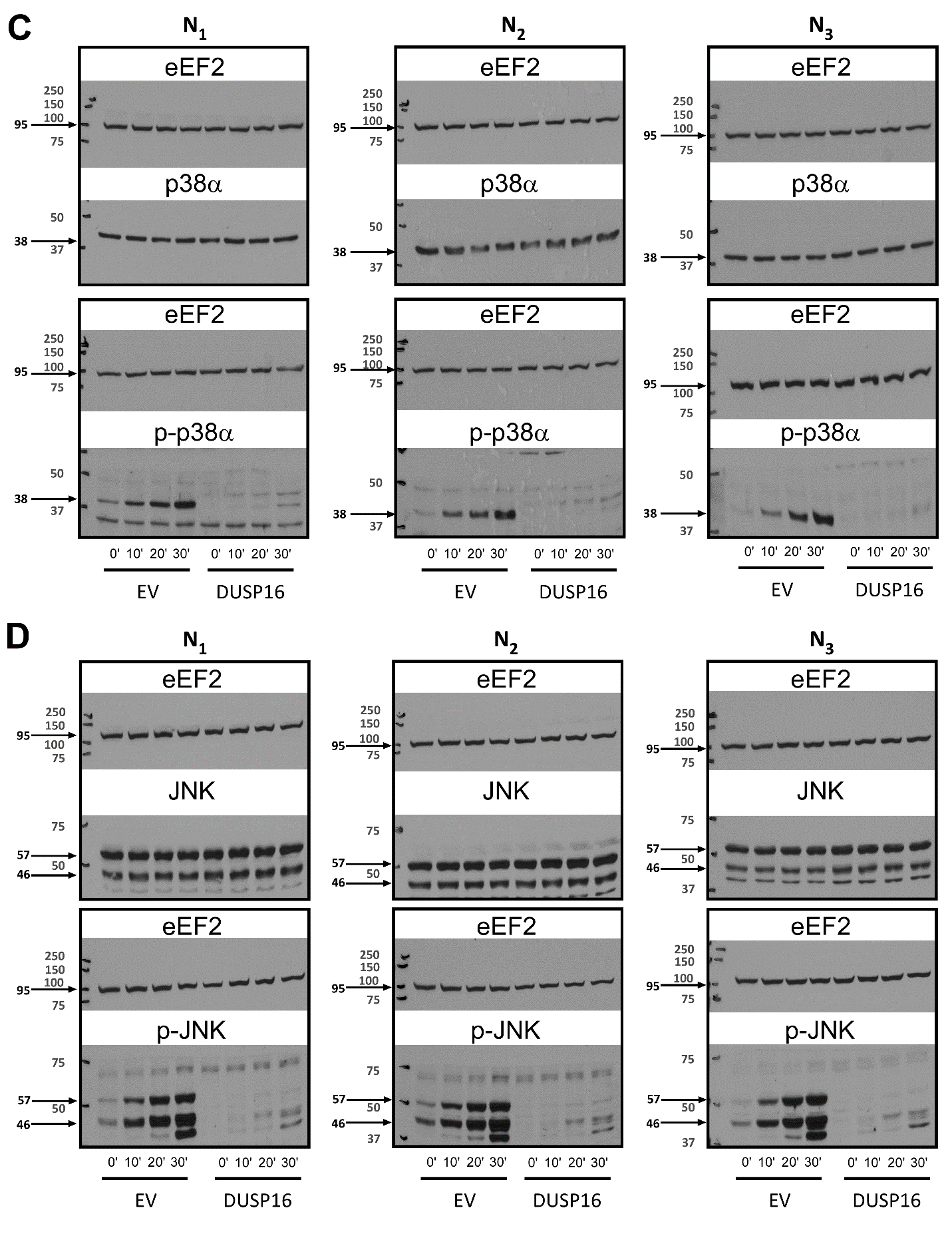

**Fig. S12: Western blots analysis (full immunoblots) of the impact of DUSP16 ORF on MAPKs phosphorylation levels in HCT-15 ((all replicates used to generate Fig.S10A&B)**

**(A)** **Impact of DUSP16 on steady-state MAPKs phosphorylation levels**. Exponentially growing HCT-15 transduced with a lentiviral pLVX-EF1a-IRES-PURO/eGFP expression plasmid, either empty (empty vector, EV) or containing the codon-optimized DUSP16 ORF (DUSP16), were grown to 80% confluence and whole cell lysates were harvested for the evaluation of the native and phosphorylated states of ERK, p38 and JNK by Western blot analysis. Whole cell lysates were run on SDS-PAGE and transferred onto nitrocellulose membranes (BioRad). The membranes were then cut in half at the level 75kDa to allow independent immunoblotting and detection of proteins of different sizes (primary test targets (MAPKs) and loading control (eEF2)); full immunoblots representing the two halves of the same membrane are shown boxed together. Region of immunoblots boxed in red are those shown in **Fig.S10A**. Molecular weight markers were run on each gel and are marked on images with corresponding sizes on the left; expected molecular weights of target proteins are indicated with arrows. Following detection, immunoblots were scanned into JPEG files and expression levels were determined using the Image J software (Fiji). Levels of each MAPK or p-MAPK were first corrected with their corresponding eEF2 loading controls, and then the level of phosphorylation of each MAPK, expressed as the ratio of phosphorylated MAPK over total MAPK, was calculated. The results are summarized in **Fig.S10A** combining the replicates (n=2 for EV, n=3 for DUSP16).

**(B-C-D)** **Impact of DUSP16 on the induction of MAPKs phosphorylation following serum starvation**. Using the same cell models as above, exponentially growing cells were first serum-starved for 24 hours (McCoy's 5A (Modified) medium, 0% FBS). The cells were then stimulated with 10 ng/ml TNF-α for different times to induce MAPK phosphorylation and whole cell lysates harvested for the evaluation of the native and phosphorylated states of MAPKs by Western blot analysis (as described above); immunoblots for each MAPK and p-MAPK across three replicates are shown separately for **(B)** ERK, **(C)** p38, and **(D)** JNK detection. Levels of each MAPK or p-MAPK were first corrected with their corresponding loading controls, and then the level of phosphorylation of each MAPK, expressed as the ratio of phosphorylated MAPK over total MAPK, was calculated. The induction of phosphorylation at different time points is shown in **Fig.S10B** (combining the three replicates) for the cells containing either empty vector (EV (+)) and the DUSP16 ORF stimulated with Doxycycline (DUSP16 (+)).

**BOX 1 – Genes in Cluster 1**

**IFIH1 (interferon Induced with Helicase C Domain 1; OMIM*** **606951).**

**Genetics:** The genomic region containing IFIH1 was first identified as an IBD risk locus in the 2012 IIBDGC study (rs2111485, P< 1.93 x 10^-8^) (1), where IFIH1 was suggested as the best functional candidate of six genes in this locus (1). The IIBDGC fine-mapping study, subsequently identified a single non-synonymous variant in this gene (rs35667974, I923V) associated to UC risk, confirming this as the most likely causal gene in the region. (2) IFIH1 has also been identified in association studies of multiple immune related diseases, including T1D where a protective effect was reported (3). Resequencing efforts in T1D have identified 4 infrequent/rare non-synonymous coding variants in IFIH1 showing significant protective effects (4). These included the I923V C-terminal domain (CTD) mutation (rs35667974), as well as a non-sense variant E627ter (rs35744605, leading to a truncation of part of the helicase domains and the CTD) and 2 splicing mutants (rs35337543, at splice donor site position +1 in intron 8 leading to an in-frame deletion within the first helicase domain and rs35732034, at splice donor site at position +1 in intron 14 leading to a premature termination with CTD deletion). Our targeted sequencing of the exons of protein-coding genes in IBD-associated regions (5) provided evidence for association to UC for three of these T1D variants, albeit conferring risk rather that protection (6). Specifically, in addition to the I923V mutation, both splice donor mutations identified in T1D are also associated with increased risk in IBD (rs35732034, MAF=0.8%, OR=1.3, *P*=0.002; rs35337543, MAF=1%, OR=1.3, *P*=0.003).

**Function:** The IFIH1 gene encodes the MDA5 protein, which is a pattern recognition receptor (PRP) that functions as a receptor for viral RNA. Upon binding of RNA, the CARD domains of MDA5 interact with mitochondrial antiviral signalling protein (MAVS), leading to the activation of IRF3/7 and the transcription of type I interferon stimulated genes (ISGs). These ISGs include molecules involved in inducing a cellular antimicrobial state in infected and neighbouring cells, modulating immune response, and activating the adaptive immune system (7).

**SBNO2 (Strawberry notch homologue 2; OMIM*615729).**

**Genetics:** The genomic region containing SBNO2 was first identified in a 2008 GWAS in CD (rs4807569, P< 2.12 x 10^-9^) (8), and confirmed as a CD locus in the 2012 IIBDGC study (rs2024092, P< 8.26 x 10^-22^) where SBNO2 was one of 22 genes in this locus (1). The IIBDGC fine-mapping study of this region subsequently identified a primary signal for CD with three SNPs located in three separate introns of the SBNO2 gene and all within 653 bp of each other (2). While these common variants do not appear to localize within known regulatory sequences (e.g. TFBS), their tight clustering within a single gene supports SBNO2 as the best candidate causal gene. Furthermore, a secondary association signal was detected in CD for this region for 10 SNPs located over a region of less than 11kb upstream of SBNO2, with rs72977562 located within a gut enhancer element (H3K27ac).

**Function:** The SBNO2 gene encodes a DExD/H helicase family corepressor protein known to do act as a transcriptional regulator. Expression profiling of SBNO2 indicates that SBNO2 is expressed across a wide range of tissues and immune cells, and whose expression is up-regulated by hyper-IL-6 (IL-6 + soluble IL-6 receptor) and by IL10 (9, 10). While there is limited information on the function of this gene, it has been reported that it contributes to the downstream anti-inflammatory effects of IL-10 (9) and may play a variety of roles in sepsis (11), bone homeostasis (12) and inflammatory response in the central nervous system (10).

**NFKB1 (Nuclear Factor Kappa B Subunit 1; OMIM*164011).**

**Genetics:** The genomic region containing NFKB1 was first identified in the 2012 IIBDGC study with strongest association to UC (rs3774959, P< 3.66 x 10^-12^) (1). This region was not part of the IIBDGC Fine Mapping project. While the original study identified a region containing five gene candidates, our LD-based definition of the associated region used in the current study identifies NFKB1 as the only candidate gene in this locus.

**Function:** NFKB1 is part of a family of Rel-like domain containing protein that also includes RELA/p65, RELB, NFKB2/p52. Members of this family form the NF-kappa-B transcription factor that is made-up of homo- and heterodimeric complexes with different affinities and specificities for DNA binding sites and that can act either as transcriptional activators or repressors. Thus the NF-kappa-B transcription factor can have a pleiotropic effect on cells, impacting on a wide range of biological functions including inflammation and immunity. NF-kappa-B is involved in responses to cytokines and is a key regulator of the innate immune response to bacteria. NF-kappa-B activity has been shown to be induced in a many inflammatory diseases, including IBD, where it is believed to stimulate pro-inflammatory cytokines and thus influence inflammation.

**NOD2 (nucleotide-binding oligomerization domain protein 2; OMIM*605956).**

**Genetics:** The genomic region containing NOD2 was one of the first genomic regions (IBD1 region) to be linked to CD. Fine mapping of this region, using a candidate gene approach, identified association to CD for 3 rare non-synonymous coding variants in NOD2 (R702W, G908R, and fs1007insC) (13, 14). These variants have, over the years, been reproducibly associated to CD across multiple association studies and were also shown to be strongly associated with Crohn's disease location as well as age at diagnosis. Targeted sequencing of the exons of protein-coding genes in IBD-associated regions provided evidence for association to CD for an additional six rare NOD2 variants (15). The IIBDGC fine-mapping study of this region identified multiple independent association signals to CD in this region, the majority of which included either previously described or novel rare non-synonymous coding variants in NOD2 (16). While our LD-based definition of IBD-associated regions identifies three potential candidate genes, given that NOD2 an accepted causal gene it was the only candidate gene from this region included in the current study

**Function:** NOD2 is a cytosolic pattern recognition receptors (PRRs) that belongs to the intracellular NOD-like receptor family. Activation of NOD2 by MDP (a component of bacterial cell wall peptidoglycan) triggers innate proinflammatory and anti-bacterial immune response pathways via the activation of NFkB signalling.

**ZFP36L1 (ZFP36 Ring Finger Protein Like 1; OMIM*601064).**

**Genetics:** The genomic region containing ZFP36L1 was first identified in the 2010 IIBDGC meta-analysis of CD GWAS (rs4902642, P< 1.60 x 10^-10^) (17), and defined as an IBD locus in the 2012 IIBDGC study (rs194749, P< 2.70 x 10^-10^) (1). This region was not part of the IIBDGC Fine Mapping project. While the original study identified a region containing five gene candidates, based on our definition of the associated region used in the current study only ZFP36L1 and ACTN1 were identified as candidate genes in the locus.

**Function:** ZFP36L1, is part of a family of RNA binding proteins (RBP) that bind A/U rich elements (ARE) in the 3’ untranslated region (3’UTR) of mRNAs and promotes of RNA decay. Most reports studying the role of this gene family have focused on immune cells, and have proposed a role for ZFP36 in regulating pro-inflammatory responses (through the regulation of RNA levels of inflammatory cytokines) and for ZFP36L1/L2 in T- and B-lymphocyte development. Of relevance to the current study, a recent report using a non-immune model (Hela cells) has proposed that ZFP36 could regulate gene targets at the transcriptional level, rather than RNA stability, and has shown that ZFP36 overexpression leads to an increased expression of genes involved in Type I interferon signalling and antiviral response, an effect similar to the one observed for ZFP36L1 in our model (18). The other candidate gene in this locus, ACTN1, encodes for actinin alpha 1 a ubiquitously expressed structural protein involved in cross-linking F-actin to intracellular structures. Given its role as a structural protein, it is not surprising that ACTN1 did not impact significantly on the transcriptome (7 HITS). Based on our screen, ZFP36L1 is the best functional candidate in this region.

**IRF1** **(Interferon Regulatory Factor 1; OMIM*147575).**

**Genetics:** The genomic region containing IRF1 was one of the first genomic regions (IBD5 region) to be associated to CD. While this region has, over the years, been reproducibly associated to CD across most association studies, there have also been some reports of association to UC as well. The 2012 IIBDGC study has confirmed the association of this locus to IBD, with 19 potential candidate genes identified within this region. The IIBDGC fine-mapping study of this region subsequently identified a primary signal for IBD with eight SNPs located within a 35 Kb intergenic region between IRF1 and SLC22A5, four of which impacting on gut (H3K27ac) and/or immune (H3K4me1) enhancer element (2). The tight clustering between IRF1 and SLC22A5 supports these two genes as candidate causal genes for this region.

**Function:** IRF1 is transcriptional regulator involved in activating both innate and acquired immune responses to viral and bacterial infection through the regulation of interferon and ISGs, as well as several genes involved in immune response through binding to interferon-stimulated response elements (ISRE) in their promoters. IRF1 also plays a role in the maturation and activation of different immune cell types. SLC22A5 is a member of the Solute Carrier Family 22 that encodes for OCTN2 a sodium-dependent high affinity carnitine transporter. Both SLC22A5, and its neighboring gene SLC22A4, have previously been proposed as causal genes in this genomic region based on the identification of IBD associated variants either in proximal promoter region (SLC22A5, -207C>G) or causing a missense substitution (SLC22A4, L503F).

**GIGYF1 (GRB10 Interacting GYF Protein 1; OMIM*** **612064).**

**Genetics:** The genomic region containing GIGYF1 was first identified in the 2012 IIBDGC study with strongest association to IBD (rs1734907, P< 1.67 x 10^-13^) (1). This region was not part of the IIBDGC Fine Mapping project. While the original study identified a region containing 22 gene candidates, our LD-based definition of the associated region used in the current study identifies 4 potential candidate genes in this locus: GIGYF1, ACTL6B, GNB2 and POP7.

**Function:** GIGYF1, a GYF domain-containing scaffold protein, has been proposed to regulate signalling through insulin receptors via its interaction with the adaptor protein GRB10. GIGYF1 can also promote translational repression of specific targets through its interaction with mRNA-binding proteins. Interestingly, it has been shown that one of these mRNA-binding proteins is ZFP36 family member TTP (see above, ZFP36L1). The 3 other gene candidates in this genomic region (ACTL6B, an actin-like protein; GNB2, a G-protein subunit; POP7, a component of ribonuclease P) did not impact significantly on the transcriptome (1 HIT, 5 HITS and 3 HITS respectively) and therefore it is impossible to assign them a role in epithelial tissues. Based on our screen, GIGYF1 is the best functional candidate in this region.

**OTUD3 (OTU Deubiquitinase 3; OMIM*** **611758).**

**Genetics:** The genomic region containing OTUD3 was first identified in a 2009 GWAS in UC (rs6426833, P< 5.1 x 10^-13^) (19) with at least two independent association signals detected, and confirmed as a UC locus in the 2012 IIBDGC study (rs6426833, P< 2.39 x 10^-68^) where OTUD3 was one of nine genes in this locus (1). The IIBDGC fine-mapping study subsequently identified three independent association signals to UC in this region (2), two of which containing SNPs located within a gut enhancer element (H3K27ac) near RNF186, and narrowed the list of potential candidate genes in the locus to three (TMCO4, RNF186 and OTUD3). Our targeted sequencing of the exons of protein-coding genes in IBD-associated regions provided evidence for association to UC for two rare RNF186 variants: a non-synonymous coding risk variant A64T (rs41264113, P< 8.69 x 10^-4^) (20) and a protective non-sense variant R179ter (rs36095412, P< 6.89 x 10^-7^) (5). Our LD-based definition of the associated region used in the current study identifies 2 potential candidate genes in this locus: RNF186 and OTUD3.

**Function:** OTUD3 is a deubiquitinase that can de-polyubiquitylate the tumour suppressor PTEN, leading increased stability and levels of PTEN. It has recently been shown that PTEN plays a critical role in antiviral innate immunity through the nuclear import and activation of IRF3, a transcription factor involved in IFN-b production (21) (22), and based on our results from OTUD3 may play a similar role in our intestinal epithelial cell model. The other gene in this locus, RNF186 is a ring finger E3 ligase, localized at the endoplasmic reticulum (ER), that plays a role in regulating ER-stress induced apoptosis (23). Based on our previous genetic data for RNF186 and on the shared impact of OTUD3 with other IBD-causal gene from cluster 1 in the current screen, both RNF186 and OTUD3 may still be considered strong functional gene candidates in this region.

**AIRE (Autoimmune Regulator; OMIM*** **607358).**

**Genetics:** The genomic region containing AIRE was first identified a 2008 GWAS in CD (rs762421, P< 1.41 x 10^-9^) (8), and defined as an IBD locus in the 2012 IIBDGC study (rs7282490, P< 2.35 x 10^-26^) (1) where AIRE was one of ten candidate genes in the locus. This region was not part of the IIBDGC Fine Mapping project. Based on our definition of the associated region used in the current study C21orf33, ICOSLG and AIRE were identified as candidate genes in the locus.

**Function:** AIRE is a transcriptional regulator that plays a role in promoting self-tolerance through the expression of tissue restricted antigens in the thymus. AIRE has recently been reported to also play a role in the transcription of genes associated with systemic autoimmune disease as well as interferon-γ regulated genes in activated fibroblast-like synoviocytes in RA. In addition, AIRE promoted the production and secretion of chemokines associated with disease activity in RA (24). ICOSLG being a costimulator ligand for the T-cell specific receptor ICOS was not included in our screen. C21orf33, also known as GATD3A encodes a potential mitochondrial glutamine amidotransferase; lentiviral overexpression of this ORF did not achieve detectable levels in HT29 and therefore C21ORF33 was rejected from our screen.

**PITX1 (Paired Like Homeodomain 1; OMIM*602149).**

**Genetics:** The genomic region containing PITX1 was first identified in the 2011 IIBDGC meta-analysis of UC GWAS (rs254560, P< 1.25 x 10^-9^) (17), confirmed as a UC locus in the 2012 IIBDGC study (rs254560, P< 2.55 x 10^-9^) where PITX1 was one of six candidate genes in this locus (1). This region was not part of the IIBDGC Fine Mapping project. Based on our definition of the associated region used in the current study, only PITX1 and H2AFY were identified as candidate genes in the locus.

**Function:** PITX1 is a bicoid-related homeobox transcription factor of the RIEG/PITX family involved in organ and limb development and patterning, as well as left-right asymmetry. PITX1 was originally identified to play a role in the activation of pituitary transcription of a pro-opiomelanocortin gene. It has been reported that PITX1 can suppress the interferon α (INF-α) promoter via interaction with IRF3 and IRF7 proteins (25), while it was suggested that PITX1 played a role for upregulating of IFN-regulated genes in cutaneous lupus erythematosus lesions (26). More recently, reduced PITX1 levels in CRCs have been suggested as a marker for poor prognosis. The other gene in the region, H2AFY, encodes a histone protein; H2AFY had practically no impact on transcriptome (8 HITS).

**FOS (Fos Proto-Oncogene, AP-1 Transcription Factor Subunit; OMIM*** **164810).**

**Genetics:** The genomic region containing FOS was first identified in the 2012 IIBDGC study with strongest association to IBD (rs4899554, P< 2.71 x 10^-8^) (1). This region was not part of the IIBDGC Fine Mapping project. While the original study identified a region containing eight gene candidates, based on our definition of the associated region used in the current study, we identify 2 potential candidate genes in this locus: FOS and TMED10.

**Function:** FOS is one of four FOS family members, along with FOSB, FOSL1, and FOSL2, which can heterodimerize with members of the JUN family to form the AP-1 transcription factor. FOS has been proposed to regulate several cellular processes such as proliferation, apoptosis, differentiation and transformation. AP-1 has been shown to play a role in TGF-beta signalling. Two other members of the FOS family, FOSL1 and FOSL2, have also been identified in IBD-associated regions. A role for FOS in the regulation of interferon-beta has previously been proposed (27). The other gene in the region, TMED10, a member of the EMP24/GP25L/p24 family encodes a type I transmembrane protein of the plasma membrane involved in vesicular trafficking; TMED10 had practically no impact on transcriptome (5 HITS).

**FOXO1 (Forkhead Box O1; OMIM*** **136533).**

**Genetics:** The genomic region containing FOXO1 was first identified in the 2011 IIBDGC meta-analysis of UC GWAS (rs941823, P< 3.82 x 10^-12^) (17), and defined as an IBD locus in the 2012 IIBDGC study (rs941823, P< 2.07 x 10^-14^) (1) where FOXO1 was one of three candidate gene candidates in the locus. This region was not part of the IIBDGC Fine Mapping project. Based on our definition of the associated region used in the current study, FOXO1 was the only candidate gene in the locus.

**Function:** FOXO1 is a transcription factor of the forkhead family involved in metabolic homeostasis and adipocyte differentiation. FOXO1 transcriptional activity is inhibited via phosphorylation by AKT in response to insulin. It has been reported that FOXO1 can suppress virus-triggered IFN-β induction and cellular antiviral response via the interaction with IRF3 promoting the polyubiquitination and degradation of IRF3 in the cytosol (28).

**BOX 2 – Genes in Cluster 2**

**KSR1 (Kinase Suppressor of Ras 1; OMIM*601132).**

**Genetics:** The genomic region containing KSR1 was first identified as a CD locus in the 2012 IIBDGC study (rs2945412, P< 8.68 x 10^-17^) (1). The IIBDGC fine-mapping study, subsequently mapped this association signal to CD for seven common variants across 25kb, all within the KSR1 gene (2), highlighting KSR1 as the most likely causal gene in the region.

**Function:** KSR1 is a very good candidate gene for its locus, based on published functional results that support its role in protecting against intestinal inflammation. Specifically, Polk and colleagues observed that KSR1 is activated in inflamed mucosa and using KSR1-deficient mice demonstrated that KSR1 protects intestinal epithelium from cytokine-mediated apoptosis during inflammation (29). KSR1 has been shown to be a scaffold protein with the ability to bind Raf-1, MEK, and ERK. Scaffold proteins, like KSR1, play a critical role in ERK regulation by sequestering/compartmentalizing ERK signalling to specific cellular localisation and therefore regulate signal strength, spatial distribution and specificity. KSR1 has been shown to increase EGF-induced cytoplasmic ERK signaling via interaction with the EGF-EGFR signalling complex at the plasma membrane (30).

**DUSP16 (Dual-specificity phosphatase 16; OMIM*** **607175).**

**Genetics:** The genomic region containing DUSP16 was first identified in the 2012 IIBDGC study, with strongest association to UC (rs11612508, P< 1.06 x 10^-8^) (1). This region was not included in the IIBDGC Fine Mapping project. While the original study identified a region containing nine gene candidates, our LD-based definition of the associated region used in the current study identifies DUSP16 as the only candidate gene in this locus.

**Function:** DUSP16 is a member of the dual-specificity phosphatases (DUSPs) family of proteins that can dephosphorylate tyrosine and serine/threonine residues of specific substrates. More specifically, DUSP16 is a member of a subgroup of these, dubbed typical DUSPs or mitogen-activated protein kinase (MAPK) phosphatases (MKPs), possessing both a nuclear localization signal (NLS) and a conserved dual specificity phosphatase catalytic domain (DSP) which targets MAPKs and plays a central role in negatively regulating their activity. In addition to these domains, DUSP16 (aka MKP-7) also possesses a nuclear export signal (NES) which shuttles nuclear DUSP16 to the cytoplasm. While DUSP16 can bind all MAPK with equal affinity, it suppresses MAPKs activity with the order of selectivity, JNK ≫ p38 > ERK and is defined as a JNK-specific phosphatase *in vivo* (31). Because of its NES, DUSP16/MKP-7 has been shown to shuttle its MAPK substrates from the nucleus to the cytoplasm. DUSP16 was also shown to increase and prolong EGF-stimulated cytoplasmic ERK phosphorylation, through its ability to bind and retain pERK1/2 in the cytoplasm, while inhibiting its nuclear effect on expression of ERK target genes (32). An alternate mechanism was proposed for this increase in EGF-stimulated cytoplasmic ERK phosphorylation, suggesting that DUSP16 inhibits p38 and thus p38-driven inhibition of ERK activity (33). This illustrates the ability for DUSP16 to act as either a positive or negative regulator of cytoplasmic or nuclear ERK respectively, while still retaining its ability to directly dephosphorylate its preferred MAPK target(s).

**CDC37 (Cell Division Cycle 37; OMIM*** **605065)**

**Genetics:** The genomic region containing CDC37 was first identified in the 2010 IIBDGC meta-analysis of CD GWAS (rs12720356, P< 1.40 x 10^-12^) (17), and further defined as an IBD locus in the 2012 IIBDGC study (rs11879191, P< 2.04 x 10^-18^) (1) where CDC37 was one of 28 gene candidates in the region. The IIBDGC fine-mapping study subsequently identified three independent association signals, including a non-synonymous variant in TYK2 and a variant within a conserved transcription-factor binding site near TYK2 (2), pinpointing TYK2 as the main candidate causal gene in this region. Our LD-based definition of the associated region used in the current study identifies 3 potential candidate genes in this locus: TYK2, CDC37 and PDE4A.

**Function:** While TYK2, which is a member of the JAK tyrosine kinase family, is a strong functional candidate for CD in this locus, based on its function in intracellular signal transduction from cytokine receptors and the presence of a non-synonymous coding variant, our study highlights CDC37 as another potential functional candidate in this locus based on shared function with 4 other IBD candidate genes in cluster 2. CDC37 is part of the Hsp90/Cdc37 molecular chaperone system involved in proper folding and maturation of multiple kinases. CDC37 is a co-chaperone with a central role in connecting Hsp90 to its kinase targets (client protein), with close to 300 client proteins identified (34). CDC37 was shown to be part of a multimolecular protein complex in human embryonic kidney 293T cells, along with Hsp90 partner, where this latter protein was proposed to be essential in stabilizing KSR1 (an IBD candidate gene that is also part of cluster 2 (see above)). PDE4, the third candidate gene in the locus is a member of the cyclic nucleotide phosphodiesterase (PDE) subfamily 4 that can regulate second messenger signalling via the hydrolysis of cAMP. This gene was not evaluated in transcriptomic screen.

**TAGAP (T-Cell Activation RhoGTPase-Activating Protein; OMIM*** **609667)**

**Genetics:** The genomic locus containing TAGAP was first identified in the 2010 IIBDGC meta-analysis of CD GWAS (rs212388, P< 2.30 x 10^-11^) (17), and confirmed as a CD locus in the 2012 IIBDGC study (rs212388, P< 3.04 x 10^-14^) (1) where TAGAP was one of six gene candidates in the region. . The IIBDGC fine-mapping study subsequently narrowed down the list of potential candidate IBD genes from six to two, including TAGAP (2) and RSPH3. Based on our LD-based definition of the associated region used in the current study, TAGAP and FNDC1 were identified as candidate genes in the locus.

**Function:** TAGAP is a member of the Rho-GTPase activator family, acting as a molecular switch via the release of GTP from GTP-bound Rho. Little is known about the exact role of TAGAP in immune function, it has been found to be expressed in activated T cells and co-regulated with IL2; it is expected to play a role in T-cell activation (35). While expression of TAGAP is strongest in immune cell types and tissues, detectable expression can be seen in intestinal tissues (Table S1and protein atlas) suggesting a potential role for this gene in epithelial function. In addition to CD, TAGAP has been associated to other RA (36), Celiac (37), MS (38) and T1D, although the directionality of effect differs in depending on disease. FNDC1, the second candidate gene in the locus, a fibronectin Type III Domain Containing extracellular protein was not included in our screen.

**TTPAL (alpha tocopherol transfer protein like; no OMIM id)**

**Genetics:** The genomic locus containing TTPAL was first identified in a 2009 UC GWAS (rs6017342, P< 8.5 x 10^-17^) (39), and confirmed as a UC locus in the 2012 IIBDGC study (rs6017342, P< 1.43 x 10^-43^) (1) where TTPAL was one of 11 candidate genes. The IIBDGC fine-mapping study subsequently identified 2 independent signals to UC in the region, including a single variant signal upstream of HNF4A overlapping the H3K27ac marker in gut tissues, and narrowed down the list of potential candidate genes from 11 to six (2). Based on our definition of the associated region used in the current study, HNF4A and TTPAL were identified as the candidate genes in the locus.

**Function:** Little is known about the function of TTPAL, other than its potential in binding hydrophobic ligands such as tocopherol, and that it may have transporter activity. While it is found within cluster 2, its impact on the transcriptome is minimal (20 HITS) and does not show any strongly enriched or shared functions with other members of cluster 2. HNF4A is a much stronger functional candidate for UC in this locus based on its role as a critical transcription factor involved in the differentiation and function of intestinal epithelial cells.

**FOXD1 (Forkhead Box D1; OMIM*** **601091)**

**Genetics:** The genomic locus containing FOXD1 was first identified in the 2010 IIBDGC meta-analysis of CD GWAS (rs7702331, P< 5.90 x 10^-12^) (17), and confirmed as a CD locus in the 2012 IIBDGC study (rs7702331, P< 5.63 x 10^-10^) (1) where FOXD1 was one of four gene candidates in the region. This region was not part of the IIBDGC Fine Mapping project. Based on our definition of the associated region used in the current study, TMEM174 and FOXD1 were identified as the candidate genes in the locus.

**Function:** FOXD1 is a member of the forkhead transcription factor family; while little is known on its function; it has been involved in kidney development and may play a role in controlling inflammatory reactions and autoimmunity. FOXD1 is found within cluster 2, but its impact on the transcriptome is minimal (16 HITS) and does not show any strongly enriched or shared functions with other members of cluster 2. TMEM174, a protein of unknown function, did not show expression in intestinal tissues and was therefore not included in our study.

**SMAD7 (SMAD Family Member 7; OMIM*** **602932)**

**Genetics:** The genomic locus containing SMAD7 was first identified in the 2012 IIBDGC study, with strongest association to IBD (rs7240004, P< 1.31 x 10^-9^) (1). This region was not part of the IIBDGC Fine Mapping project. While the original study identified a region containing three gene candidates, based on our definition of the associated region used in the current study, SMAD7 and CTIF were identified as the candidate genes in the locus.

**Function:** Our study highlights SMAD7 as the potential functional candidate in this locus based on shared function with 4 other IBD candidate genes in cluster 2. SMAD7 is a nuclear protein that interacts with TGFBR1 leading to its degradation; SMAD7 is therefore an antagonist of TGF-β signaling. TGF-β signaling has a potent immune-suppressive and anti-inflammatory effect on immune cells (40). Increased mucosal SMAD7 levels in IBD patients may have an antagonistic effect on TGF-β signaling and play a role play a role disease development (41). Inhibition of SMAD7 via antisense oligonucleotide in CD has been shown to restore SMAD3 signaling and TGF-β activation, leading to suppression of inflammation (41).TGF-β signalling is also a potent inducer of epithelial-mesenchymal transition (EMT) promoting polarized epithelial cells to differentiate into motile mesenchymal cells (42). SMAD7, through its antagonistic effect on TGF-β signaling may play a role in maintaining intestinal epithelial cell function. CTIF, which is a component of the translational initiation complex and plays a role in premature termination codon recognition, was not evaluated in transcriptomic screen.

**SATB1 (Special AT-Rich Sequence Binding Protein 1; OMIM*602075)**

**Genetics:** The genomic locus containing SATB1 first identified in the 2010 IIBDGC meta-analysis of CD GWAS (rs13073817, P< 6.70 x 10^-9^) (17), and defined as an IBD locus in the 2012 IIBDGC study (rs4256159, P< 9.00 x 10^-15^) (1). This region was not part of the IIBDGC Fine Mapping project. The original study identified a region that did not contain any gene candidates; based on our definition of the associated region used in the current study, SATB1 and KCNH8 were identified as the candidate genes in the locus.

**Function:** SATB1 is a nuclear protein that binds AT-rich motifs of heterochromatin leading to chromatin remodeling and playing a role as a gene regulator. SATB1 was originally described to play a role in T-cell development and differentiation (43). More recently, increased SATB1 levels in CRC have been shown to promote EMT and metastasis, and as such have been suggested as a marker for aggressive CRC and poor prognosis (44). KCNH8, the pore-forming (alpha) subunit of the voltage-gated potassium channel subfamily 8, did not show expression in intestinal tissues and was therefore not included in our study.

**PPM1G (Protein Phosphatase, Mg2+/Mn2+ Dependent 1G; OMIM*** **605119)**

**Genetics:** The genomic locus containing PPM1G was first identified in the 2010 IIBDGC meta-analysis of CD GWAS (rs780093, P< 4.86 x 10^-11^) (17), and confirmed as a CD locus in the 2012 IIBDGC study (rs1728918, P< 9.00 x 10^-16^) (1) where PPM1G was one of 24 gene candidates in the region. This region was not part of the IIBDGC Fine Mapping project. Based on our definition of the associated region used in the current study, PPM1G and NRBP1 were identified as the only candidate genes in the locus.

**Function:** PPM1G is a member of the PP2C Ser/Thr phosphatase family that act as negative regulators of cellular stress response. PPM1G is involved in regulating spliceosome assembly (45), cell cycle progression (46) and potentially hypoxic response (through inhibition of HIF-1α levels) (47). PPM1G has a strong impact on transcriptome, with 100 HITS, and while it is identified as a member of cluster 2, functional enrichment analyses of these HITS identify terms more related to ORFs from cluster 1: TFBS analysis (IRF response elements), GO biological processes (defense response to virus, type I interferon signaling), Reactome (Interferon alpha/beta signaling). The other gene candidate from this region, NRBP1, had practically no impact on transcriptome (8 HITS). Little is known about the function of NRBP1.

**YDJC (YdjC Chitooligosaccharide Deacetylase Homolog; no OMIM id)**

**Genetics:** The genomic locus containing YDJC was first identified in the 2010 IIBDGC meta-analysis of CD GWAS (rs181359, P< 4.80 x 10^-16^) (17), and defined as an IBD locus in the 2012 IIBDGC study (rs2266959, P< 1.39 x 10^-16^) (1) where YDJC was identified as one of 13 candidate genes in the region. The IIBDGC fine-mapping study subsequently narrowed down the list of potential candidate IBD genes from 13 to five (2). Based on our definition of the associated region used in the current study, UBE2L3 and YDJC were identified as candidate genes in the locus.

**Function:** Little is known about the function of YDJC, other than its probable role in the deacetylation of acetylated carbohydrates in oligosaccharide degradation. It has also been reported that YDJC may be involved in lung cancer progression and metastasis through keratin 8 reorganization in lung cancer cells. UBE2L3, a member of the E2 ubiquitin-conjugating family, was not evaluated in transcriptomic screen.
